# Stochastic asymmetric repartition of lytic machinery in dividing CD8^+^ T cells generates heterogeneous killing behavior

**DOI:** 10.1101/2020.09.08.287029

**Authors:** Fanny Lafouresse, Romain Jugele, Sabina Müller, Marine Doineau, Valérie Duplan-Eche, Eric Espinosa, Marie-Pierre Puissegur, Sébastien Gadat, Salvatore Valitutti

## Abstract

Cytotoxic immune cells are endowed with a high degree of heterogeneity in their lytic function, but how this heterogeneity is generated is still an open question. We therefore investigated if human CD8^+^ T cells could segregate their lytic components during telophase, using imaging flow cytometry, confocal microscopy and live cell imaging. We show that CD107a^+^-intracellular vesicles, perforin and granzyme B unevenly segregate in a constant fraction of telophasic cells during each division round. Mathematical modeling posits that unequal lytic molecule inheritance by daughter cells results from the random distribution of lytic granules on the two sides of the cleavage furrow. Finally, we establish that the level of lytic compartment in individual CTL dictates CTL killing capacity. Together, our results show the stochastic asymmetric distribution of effector molecules in dividing CD8^+^ T cells. They propose uneven mitotic repartition of pre-packaged lytic components as a mechanism generating non-hereditary functional heterogeneity in CTL.

## Introduction

Heterogeneity and plasticity of lymphocyte function are key components of successful adaptive immune responses. Accordingly, several studies put forth the notion that individual mouse and human lymphocytes exhibit high degrees of heterogeneity in both their phenotypic and functional characteristics (Beuneu et al., 2010; Buchholz et al., 2016, 2013; Ganesan et al., 2017; Kumar et al., 2018; Lemaitre et al., 2013; Newell et al., 2012). Functional heterogeneity is not limited to cell differentiation and acquisition of phenotypic and functional characteristics, but also involves late steps of immune cell responses such as CD8^+^ cytotoxic T lymphocyte (CTL)- and natural killer (NK) cell-mediated cytotoxicity (Guldevall et al., 2016; Halle et al., 2016). Accordingly, we have previously shown that human CTL belonging to the same clonal population exhibit heterogeneity in their lytic function during sustained interaction with target cells (Vasconcelos et al., 2015). While, some CTL kill a limited number of target cells, others emerge as super-killer cells.

One proposed mechanism of functional heterogeneity generation in T lymphocytes is asymmetric cell division (ACD). ACD is a key mechanism to generate cell heterogeneity in biology. It plays a crucial role in embryogenesis by allowing the formation of two distinct cells from a single mother cell (Dewey et al., 2015; Knoblich, 2008). In immunology, ACD has been proposed as a process allowing mouse naive T lymphocytes to divide into short-lived effector T cells and memory T cells, after TCR-triggered division (Arsenio et al., 2015, 2014; Chang et al., 2011, 2007).

In the present work, we investigated the possibility that, in dividing human CD8^+^ T cells, heterogeneous distribution of molecules relevant for cytotoxic function into nascent daughter cells might contribute to CTL killing heterogeneity.

To address this question, we employed imaging flow cytometry, 3D confocal laser scanning microscopy, live-cell imaging and mathematical modeling to investigate whether and how lytic components might differently segregate in telophase.

Our results show that both freshly isolated human peripheral blood CD8^+^ T cells and clonal CTL exhibit a heterogeneous repartition of lytic machinery in telophase during TCR-triggered proliferation which is not part of a classical ACD process. Furthermore, we demonstrate that heterogeneous lytic compartment repartition resets at each round of CTL division and is consequently stationary but not hereditary. Finally, we show that the level of lytic granule expression in individual CTL influences their killing ability.

Together, our results unveil a mechanism of stochastic uneven repartition of pre-packaged lytic components within intracellular vesicles that generates functional plasticity during division and contributes to lytic function heterogeneity of individual cells belonging to clonal populations.

## Results

### Imaging flow cytometry reveals uneven repartition of lytic machinery in dividing human CD8^+^ T cells

To investigate the mechanisms leading to the generation of CTL exhibiting heterogeneous killing ability, we first measured the distribution of lytic machinery components in dividing human CD8^+^ T cells. Telophase is the *bona fide* cell cycle phase where unambiguous measurement of molecular repartition in nascent daughter cells is performed (Chang et al., 2007; Filby et al., 2011). Lytic granule repartition during human CD8^+^ T cell division was evaluated using imaging flow cytometry, a technique that combines the advantages of both flow cytometry and microscopy (Basiji and O’Gorman, 2015; Doan et al., 2018; Hritzo et al., 2018). This approach allowed us to collect and analyze a substantial number of cells and to visualize and assess the repartition of molecules of interest within individual cells that were unambiguously identified as being in telophase. Cells in telophase were identified using a computer-assisted gating strategy, on the basis of nuclear and tubulin stainings (**Figure 1-figure supplement 1**). Nuclear staining with SYTOXorange^©^ identified bi-nucleated cells with elongated shape corresponding to cells in the late steps of division (anaphase and telophase). The cells in telophase were identified (and discriminated from possible cellular doublets) on the basis of tubulin staining that allowed us to highlight their midbodies. **Figure 1-figure supplement 2A** shows how masks were applied to delimit the cells and measure the fluorescence intensity of markers of interest in the nascent daughter cells. Cells were also stained with Cell Trace Violet^©^ (CTV), a probe that labels total cell proteins. As previously reported (Filby et al., 2011), we observed that total proteins distribute in nascent daughter cells within a range of 40-60% (**Figure 1-figure supplement 2B**). In our study, CTV staining served both as a marker of cell division (allowing us to identify cells in the different division rounds (Quah and Parish, 2012)), and to define total protein repartition in telophase (Filby et al., 2011). This procedure minimized the possibility that, if some images were taken slightly on an angle, with one daughter cell slightly more in focus than the other, the markers of interest would artificially appear as asymmetric. Indeed, asymmetric distribution was defined as cells in telophase in which repartition of the marker of interest in the nascent daughter cells was beyond the 40-60 % limits observed for CTV repartition (**Figure 1-figure supplement 2B**). In addition, to further exclude the possibility of measurement artifacts, we verified individual cells by eyes and included in the analysis only cells in telophase that were on a even plane. Specificity of staining for the various markers was validated (see Material and Methods section).

In a first approach, CD8^+^ T cells freshly isolated from healthy donor blood samples were stimulated with immobilized anti-CD3/anti-CD28/ICAM-1 for 72 hours. Anti-CD3/anti-CD28/ICAM-1 stimulation resulted in activation of human CD8^+^ T cells as shown by cell proliferation and CD137 up-regulation (**Figure 1-figure supplement 3**). Repartition of the lysosomal marker CD107a was investigated in cells in telophase. As shown in **Figure 1A**, while CTV distribution ranged between 40-60% in dividing T cells, 23 % of telophasic CD8^+^ T cells exhibited an uneven distribution of CD107a^+^ vesicles overcoming the 40-60% CTV range.

**Figure 1:**
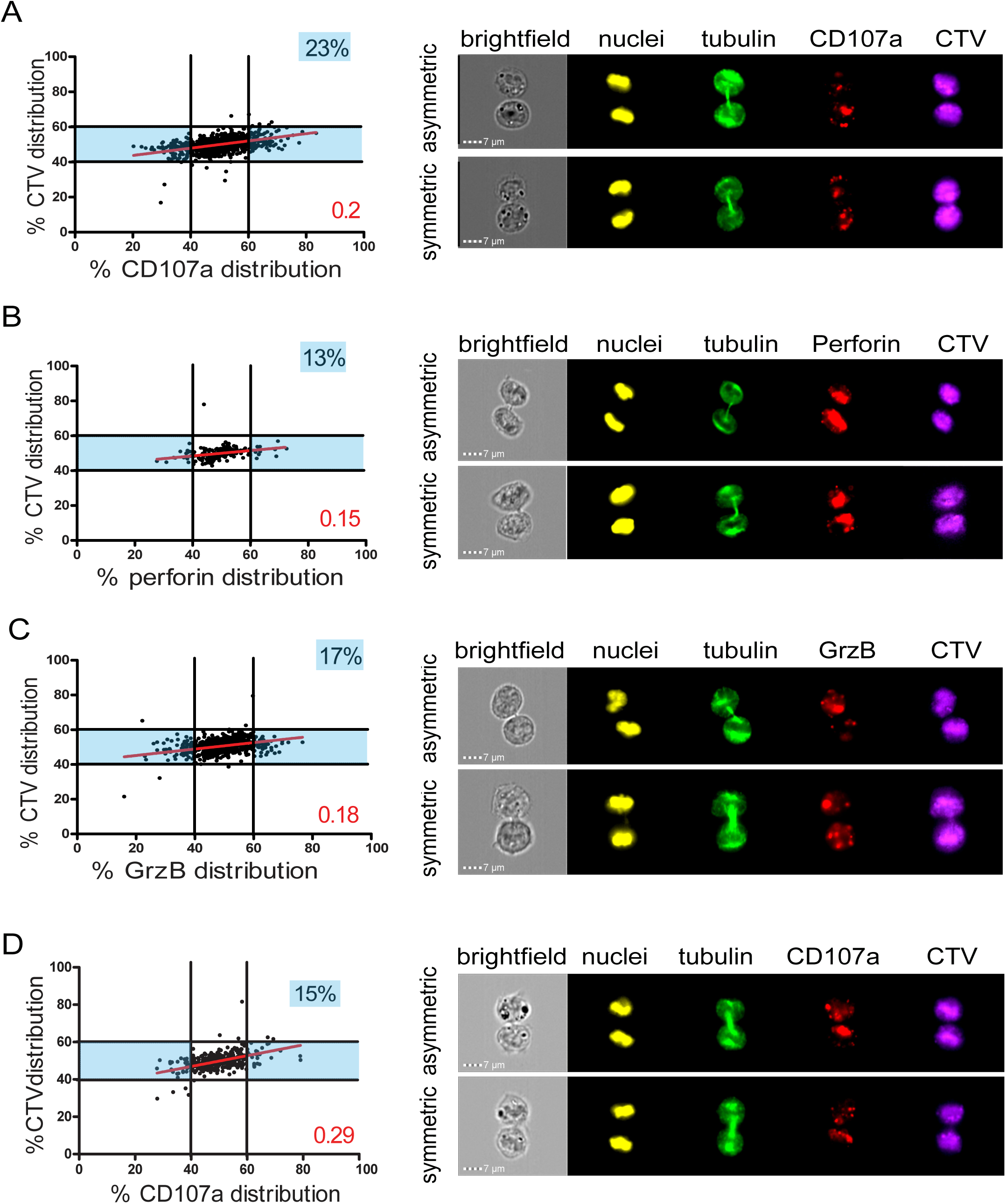
Lytic components are asymmetrically distributed in dividing CD8^+^ T cells. (**A-C**) Freshly isolated polyclonal CD8^+^ T cells or (**D**) CTL clones were stimulated by immobilized anti-CD8/anti-CD28/ICAM-1 during 72h and stained with antibodies directed against the indicated markers. Cells in telophase were identified using Imaging Flow Cytometry **(A)** Left panel: Each dot represents one nascent daughter cell. Only one of the two nascent daughter cells in telophase is plotted. The percentage of staining for CD107a in the presented cell (*x axis*) is plotted against the percentage of staining for total cell proteins (CTV, *y axis*). Asymmetric cells were defined as cells in telophase in which repartition of CD107a in the nascent daughter cells was beyond the 40-60% observed for CTV repartition (n=908 from 3 independent experiments). Right panel: example of asymmetric and symmetric cell distribution of CD107a, as detected by Imaging Flow Cytometry. **(B)** Left panel: The percentage of staining for perforin in the presented nascent daughter cell is plotted as in panel A. Asymmetric cells were defined as indicated in panel A (n=191 from 3 independent experiments). Right panel: example of asymmetric and symmetric cell distribution of perforin. **(C)** Left panel: The percentage of staining for GrzB in the presented nascent daughter cell is plotted as in panel A. Asymmetric cells were defined as indicated in panel A (n=728 from 2 independent experiments). Right panel: example of asymmetric and symmetric cell distribution of GrzB. (**D**) Left panel: The percentage of staining for CD107a is plotted as in panel A. Asymmetric cells were defined as indicated in panel A (n=352 from 3 independent experiments). Right panel: example of asymmetric and symmetric cell distribution of CD107a. Numbers highlighted in blue in the plots indicate the % of cells exhibiting asymmetric repartition of the marker of interest. Red lines indicate the global distribution of the data. Red numbers indicate the slope of the linear regression curve for marker distribution. See Figure S1, S2, S3 and S4.

**Table 1.** Results of independence Chi-square test in telophase

| Independence Chi2 test between heterogeneous cells and all cells | Test statistic ($\chi^2$) | $\chi^2_{1-\alpha, dl}$ | p-value (p) | Degree of freedom (dl) |
| --- | --- | --- | --- | --- |
| CD107a, Experiment 1, 0 division | 4.060439 | 11.07 | 0.540748 | 5 |
| CD107a, Experiment 1, 1 division | 3.565087 | 11.07 | 0.613563 | 5 |
| CD107a, Experiment 1, 2 divisions | 1.614763 | 7.815 | 0.656047 | 3 |
| CD107a, Experiment 2, 0 division |  |  |  |  |
| CD107a, Experiment 2, 1 division | 0.278928 | 7.815 | 0.963942 | 3 |
| CD107a, Experiment 2, 2 divisions | 0.413804 | 7.815 | 0.937376 | 3 |
| CD107a, Experiment 3, 0 division | 2.36867 | 15.51 | 0.967574 | 8 |
| CD107a, Experiment 3, 1 division | 2.092976 | 9.488 | 0.718663 | 4 |
| CD107a, Experiment 3, 2 divisions | 0.655225 | 9.488 | 0.956734 | 4 |

**Table 2.** Results of independence Chi-square test in telophase

| Kolmogorov-Smirnov test<br>Experiment 1 | 0 division |  | 1 division |  | 2 divisions |  |
| --- | --- | --- | --- | --- | --- | --- |
| | $D_{n,m}$ | p-value | $D_{n,m}$ | p-value | $D_{n,m}$ | p-value |
| 0 division |  |  |  |  |  |  |
| 1 division | 0.13148 | 0 |  |  |  |  |
| 2 divisions | 0.220034 | 0 | 0.116283 | 0 |  |  |

|  | 0 division | 1 division | 2 divisions |
| --- | --- | --- | --- |

| Kolmogorov-Smirnov test<br>Experiment 1 | $D_{n,m}$ | p-value | $D_{n,m}$ | p-value | $D_{n,m}$ | p-value |
| --- | --- | --- | --- | --- | --- | --- |
| 0 division |  |  |  |  |  |  |
| 1 division | 0.087873 | 0 |  |  |  |  |
| 2 divisions | 0.0891924 | 0 | 0.04634 | 0.03582 |  |  |
| 3 divisions | 0.054621 | 0.0159 | 0.067702 | 0.001185 | 0.047275 | 0.116534 |

Table 2: Results of Kolmogorov-Smirnov test on G1
| Kolmogorov-Smirnov test<br>Experiment 1 | 0 division |  | 1 division |  | 2 divisions |  |
| --- | --- | --- | --- | --- | --- | --- |
| | $D_{n,m}$ | p-value | $D_{n,m}$ | p-value | $D_{n,m}$ | p-value |
| 0 division |  |  |  |  |  |  |
| 1 division | 0.14714 | 0.002607 |  |  |  |  |
| 2 divisions | 0.209553 | 0 | 0.143594 | 0 |  |  |
| 3 divisions | 0.190642 | 0 | 0.121757 | 0 | 0.038549 | 0.3545 |

We next investigated the distribution in telophase of lytic components such as perforin and granzyme B (GrzB), molecules known to be pre-stored in lytic granules. As shown in **Figure 1B** and **C**, perforin and GrzB also unevenly segregated into the two nascent daughter cells in telophase, indicating that daughter cells received a heterogeneous quantity of lytic components.

The slope of the linear regression curve for the distribution of CD107a, perforin and GrzB as compared to CTV was close to 0.1, indicating that these 3 molecules distributed independently from total proteins.

To define whether uneven repartition of lytic components could be observed in fully differentiated cells, such as memory cells, we investigated CD107a and perforin distribution in telophase in purified human memory CD8^+^ T cells. This analysis showed that also memory CD8^+^ T cells exhibited uneven repartition of CD107a and perforin in telophase (**Figure 1-figure supplement 4**).

We next investigated whether lytic machinery asymmetric repartition could also be observed in activated CD8^+^ T cell populations composed of monoclonal cells such as antigen-specific CTL clones. To address this question, we investigated CD107a repartition in CTL undergoing cell division. For this study, we activated CTL clones using immobilized anti-CD3/anti-CD28/ICAM-1 for 72 hours. We opted for this stimulation condition since, in preparatory experiments, we observed that conjugation of CTL with cognate target cells, results (during the 72 hours culture) in the creation of cellular clumps and debris due to CTL killing activity, thus making it difficult and potentially misleading to analyze cells by image flow cytometry and conventional microscopy. As shown in **Figure 1D**, we observed that in clonal CTL undergoing cell division, 15% of the two nascent daughter cells in telophase exhibited uneven distribution of CD107a, thus confirming and extending observations obtained using CD8^+^ peripheral blood T cells. Taken together, the above results indicate that a lysosomal-associated membrane protein known to be a marker of lytic granules and effector molecules involved in CTL lytic function, unevenly segregate in 10-23 % of individual human CD8^+^ T cells undergoing division.

### Confocal laser scanning microscopy confirms uneven repartition of lytic machinery in dividing CD8^+^ T cells

Image flow cytometry allows the unambiguously identification and capture of rare events within a cell population, such as cells in telophase, albeit exhibiting a lower resolution when compared to classical imaging methods. This notion prompted us to confirm results obtained using imaging flow cytometry, with additional methods.

We therefore used 3D confocal laser scanning microscopy to measure CD107a content in telophasic CD8^+^ T cells following stimulation with immobilized anti-CD3/anti-CD28/ICAM-1. Although this approach allowed us to collect a relatively small number of cells in telophase (n=61 compared to n=908 obtained by image flow cytometry), it revealed that 27% of the CD8^+^ T cells in telophase exhibited uneven repartition of CD107a, above a 1.5 threshold (corresponding to the 40-60% range used in imaging flow cytometry experiments) (**Figure 2A**). **Figure 2B** depicts the maximum intensity projection (MIP) of a z-stack of images on which measurements of fluorescence intensity were performed (left panel) and a central z-section (right panel). The asymmetry of CD107a repartition in nascent daughter cells is better appreciated by looking at the 3D reconstructions of the dividing cell (**Video 1**).

**Figure 2:**
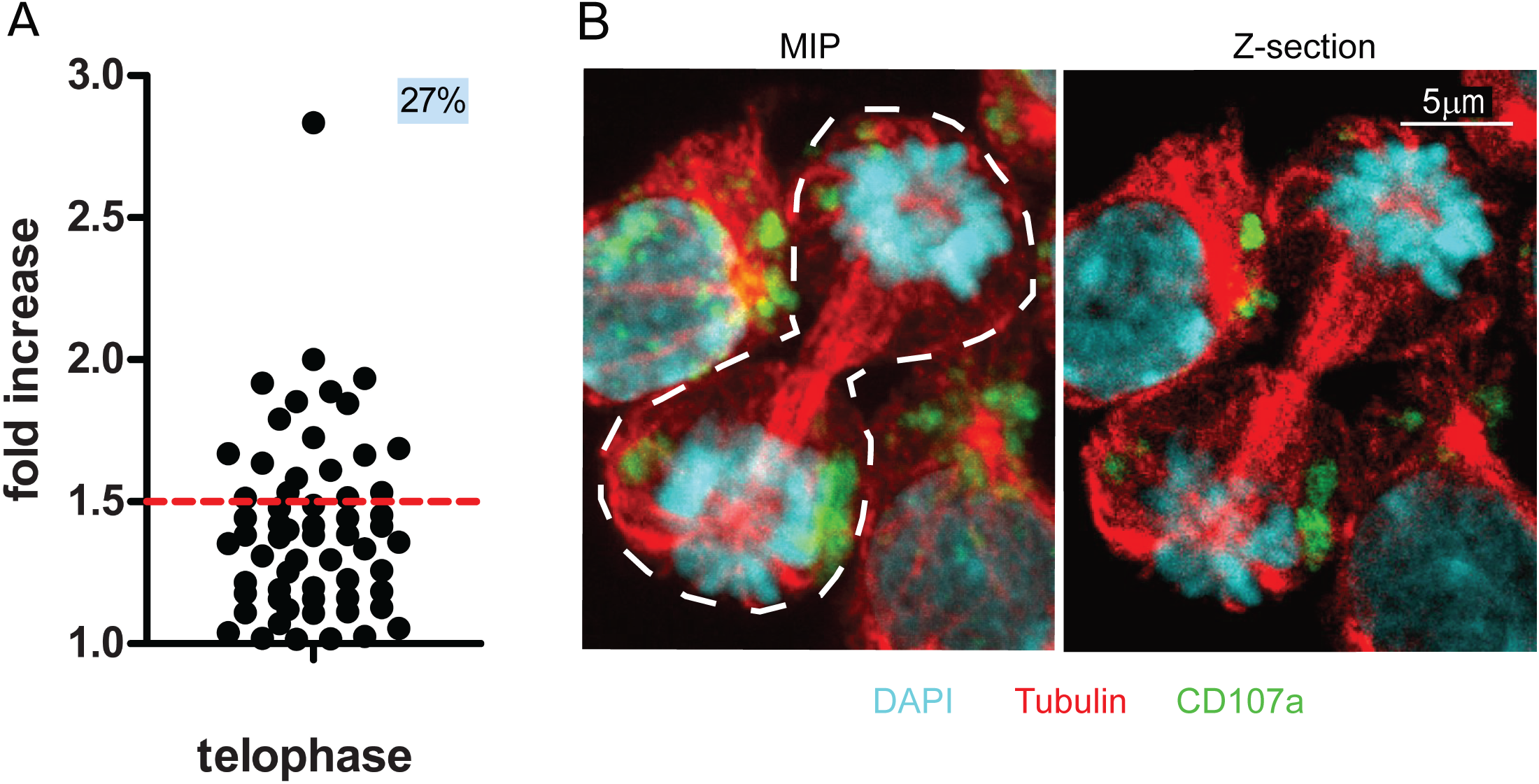
CD107a^+^ vesicles uneven segregation in telophase is confirmed by confocal laser scanning microscopy. Freshly isolated polyclonal CD8^+^ T cells were stimulated by immobilized anti-CD8/anti-CD28/ICAM-1 during 72h and stained with antibodies directed against CD107a. Cells in telophase were identified using confocal laser scanning microscopy. **(A)** Analysis of CD107a repartition in dividing cells. The fold increase of CD107a staining in the brighter nascent daughter cell as compared to the other nascent daughter cell is shown. The dotted red line indicates the limit between symmetric and asymmetric cells (1,5 fold increase, corresponding to a 60-40% variation) (n=61 from 2 independent experiments). Each dot represents one CD8^+^ T cell in telophase. **(B)** Example of an asymmetric cell in division. Green CD107a, cyan DAPI, red Tubulin. A maximum intensity projection (MIP) of a z-stack of images (left panel) and one z-section (right panel) are shown. See Video 1.

Together, the above results indicate that confocal laser scanning microscopy provides results that reinforce those we obtained using imaging flow cytometry and supports the finding that lytic granules undergo uneven repartition in ∼20% of dividing CD8^+^ T cells.

### Uneven repartition of lytic machinery is not accompanied by asymmetric segregation of fate determining transcription factors and does not require a polarity cue

The observation that lytic components were unevenly inherited in daughter cells prompted us to investigate whether this process was somehow related to mechanisms of cell fate determining ACD, a process reported to play a role in mouse naive T lymphocytes differentiation (Arsenio et al., 2015, 2014; Kaminski et al., 2016; Pham et al., 2014). Indeed, it has been reported that ACD can result in the generation of one daughter cell predisposed to become a short-lived effector cell (harboring a high level of the transcription factors T-bet and c-myc, and of GrzB) and one daughter cell predisposed to become a memory T cell (Widjaja et al., 2017). We investigated whether uneven repartition of fate determining transcription factors T-bet and c-myc (Chang et al., 2011; Verbist et al., 2016), might occur in telophase in freshly isolated peripheral blood CD8+ T cells stimulated with anti-CD3/anti-CD28/ICAM-1 for 72 hours. As shown in **Figure 3A** and **B**, both T-bet and c-myc did not unevenly segregate into the two nascent daughter cells during telophase. Moreover, the slope of the linear regression curve for the distribution of T-bet and c-myc as compared to CTV was close to 1, indicating that the repartition of these 2 molecules in telophase followed that of total proteins.

**Figure 3:**
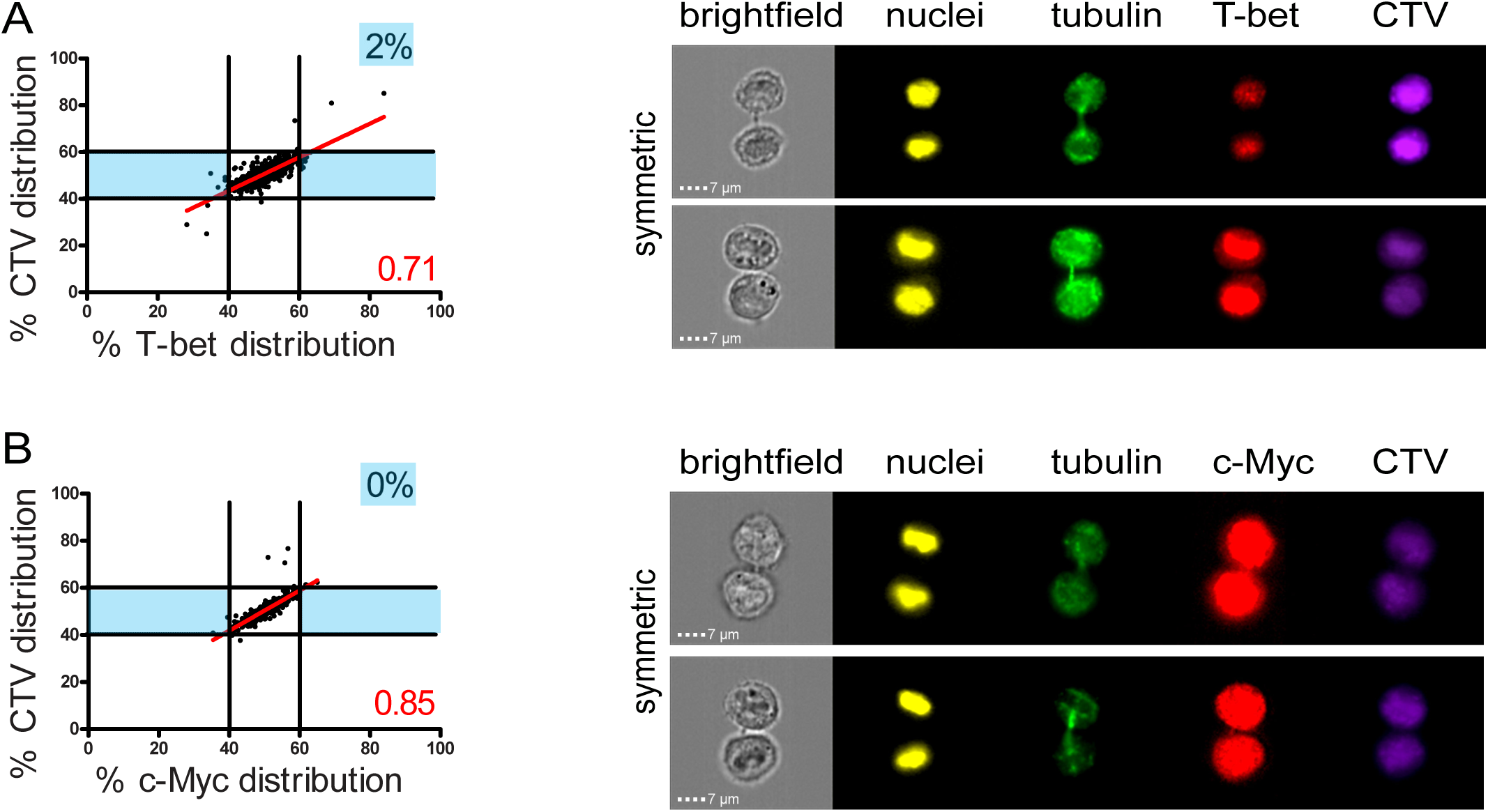
Fate determining transcription factors do not undergo uneven distribution in telophase. Freshly isolated polyclonal CD8^+^ T cells were stimulated by immobilized anti-CD8/anti-CD28/ICAM-1 during 72h and stained with antibodies directed against T-bet (**A**) or c-Myc (**B**). (**A**): T-bet analysis (n=926 from 3 independent experiments). (**B**): c-Myc analysis (n=703 from 3 independent experiments). Numbers highlighted in blue in the plots indicate the % of cells exhibiting asymmetric repartition of the marker of interest. Red lines indicate the global distribution of the data. Red numbers indicate the slope of the linear regression curve for marker distribution.

To further define whether the observed uneven repartition of lytic components was or was not related to ACD, we investigated whether uneven repartition of lytic components was dependent on a polarity cue (e.g. localized TCR stimulation) as previously described for ACD (Arsenio et al., 2015; Pham et al., 2014). **Figure 4A** and **B** shows that a polarity cue was not required to induce uneven distribution of lytic molecules, since comparable CD107a^+^ vesicles segregation was observed in peripheral blood CD8^+^ T cells stimulated by either immobilized (anti-CD3/anti-CD28/ICAM-1) or soluble (PMA + ionomycin) stimuli.

**Figure 4:**
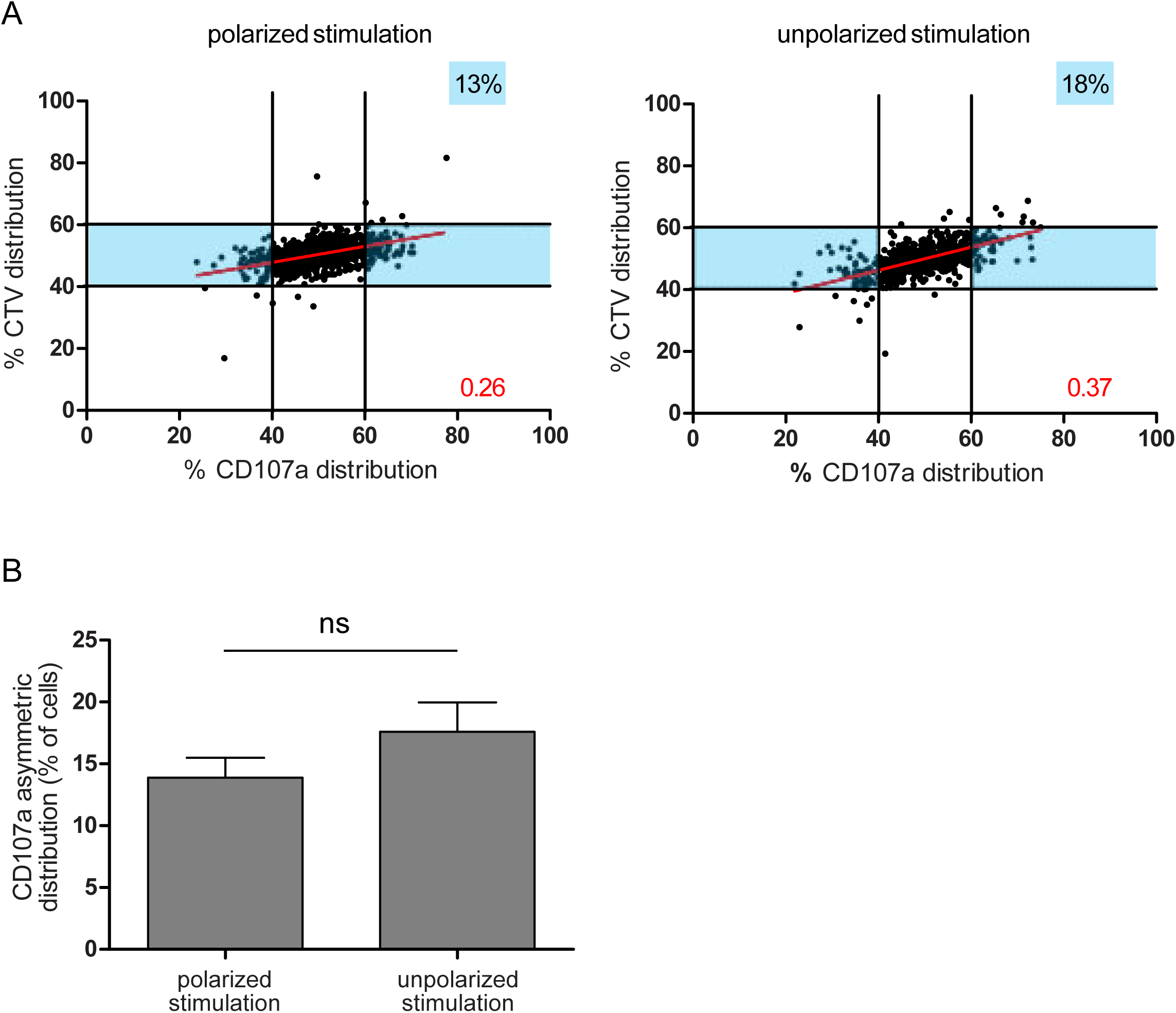
A polarity cue is not necessary for asymmetric repartition of lytic machinery. **(A)** Freshly isolated polyclonal CD8^+^ T cells were stimulated using immobilized anti-CD8/anti-CD28/ICAM-1 (left) or with PMA/ionomycin (right) during 72 hours and stained with antibodies directed against CD107a. Each dot represents one nascent daughter cell. Only one of the two nascent daughter cells in telophase that were identified by Imaging Flow Cytometry is plotted. The percentage of staining for CD107a in the presented nascent daughter cell (*x axis*) is plotted against the percentage of staining for total cell proteins (CTV, *y axis*). Asymmetric cells were defined as in Figure 1. Left: CD107a analysis when cells were stimulated with immobilized stimuli (n=1185 from 3 independent experiments). Right: CD107a analysis when cells were stimulated with PMA/ionomycin (n=644 from 3 independent experiments). Numbers highlighted in blue in the plots indicate the % of cells exhibiting asymmetric repartition of the marker of interest. Red lines indicate the global distribution of the data. Red numbers indicate the slope of the linear regression curve for CD107a distribution. **(B)** Histograms represent the mean and standard deviation of the percentage of asymmetric cells in the 3 independent experiments. No statistical difference was revealed by paired t-test.

Overall, the above results demonstrate that uneven partitioning of lytic compartment in telophase is not associated with asymmetric segregation of fate determining transcription factors. Moreover, a polarity cue is not required. All in all, the above results show that, in human CD8^+^ T cells, lytic machinery uneven repartition is not related to described mechanisms of fate determining ACD.

### Asymmetric repartition of CD107a^+^ vesicles reset at each division event and generates heterogeneous daughter cells

We next investigated whether lytic machinery uneven repartition occurred during subsequent divisions and whether this process could be involved in preserving lytic machinery heterogeneity within CD8^+^ T cell populations.

We considered the cells in the different rounds of division (identified by different peaks of CTV dilution, **Figure 1-figure supplement 3)** and analyzed CD107a repartition in telophasic cells. This analysis showed that, in all division rounds considered, a comparable percentage of cells underwent heterogeneous repartition of CD107a (**Figure 5A and B**). A complementary observation indicated that the heterogeneity process is stationary but not hereditary: e.g. a daughter cell originating from a heterogeneous division has a constant stationary probability to produce a new uneven division. We arrived to this conclusion by generating CD107a fluorescence intensity (CD107a-FI) density curves of all telophasic cells having undergone 0, 1 or 2 mitosis. Cells in telophase showing unequal CD107a-FI repartition were then plotted on these curves (**Figure 5C**). The χ^2^ statistical test showed that these cells were randomly and independently distributed on the CD107a-FI density curves, supporting the hypothesis that there is no inheritance in the decision to divide unevenly (see Materials and Methods section).

**Figure 5:**
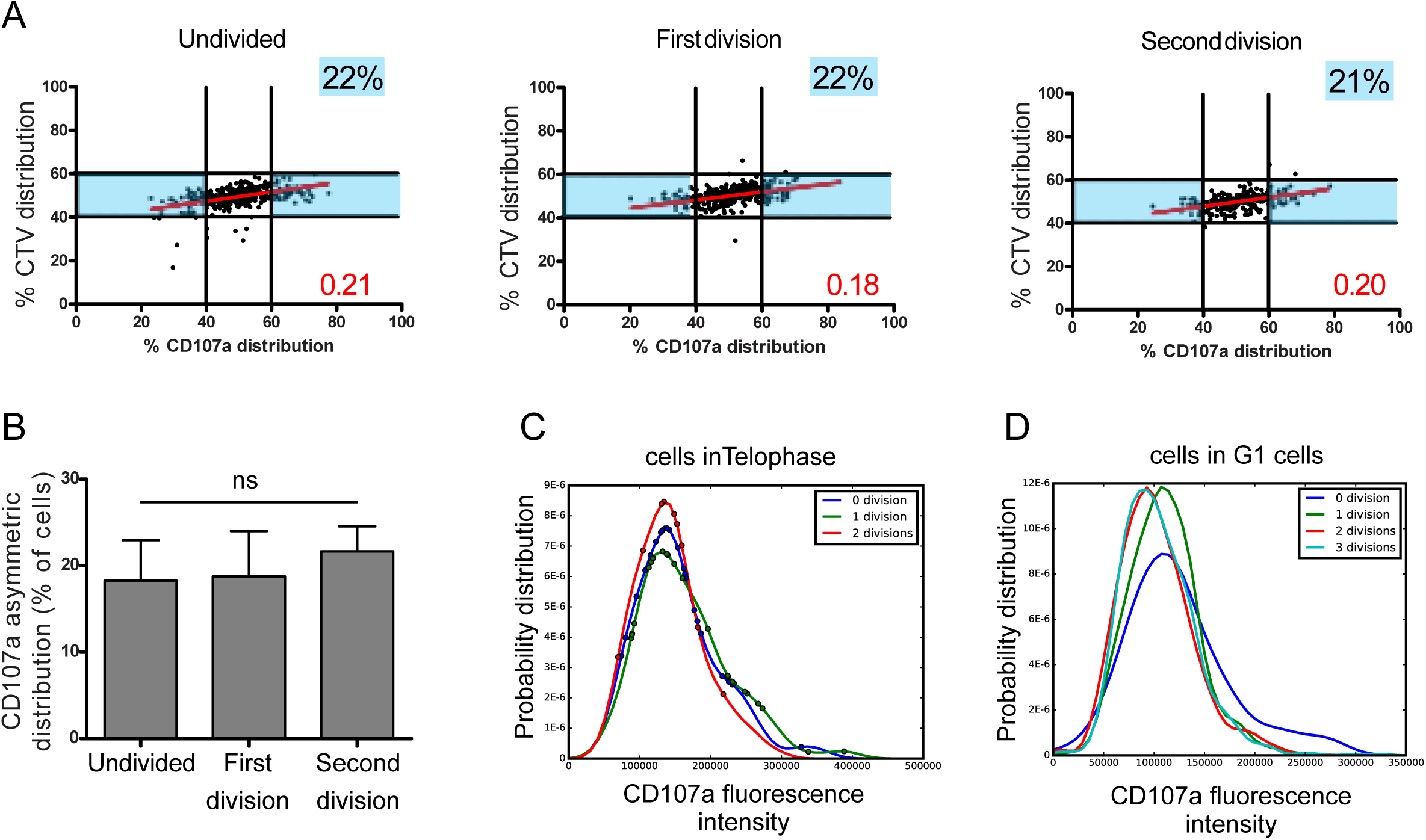
Asymmetric repartition of CD107a^+^ vesicles reset at each division event. **(A, B)** Freshly isolated polyclonal CD8^+^ T cells were stimulated using immobilized anti-CD8/anti-CD28/ICAM-1 during 72h and stained with antibodies directed against CD107a. Cells in telophase were identified by Imaging Flow Cytometry. The number of divisions accomplished and the cell cycle phase were determined on the basis of CTV and SYTOX nuclear staining. **(A)** Each dot represents one nascent daughter cell. Only one of the two nascent daughter cells in telophase that were identified by Imaging Flow Cytometry is plotted. The percentage of staining for CD107a in the presented nascent daughter cell (*x axis*) is plotted against the percentage of staining for total cell proteins (CTV, *y axis*). Asymmetric cells were defined as in Figure 1. Numbers highlighted in blue in the plots indicate the % of cells exhibiting asymmetric repartition of the marker of interest. Red lines indicate the global distribution of the data. Red numbers indicate the slope of the linear regression curve for CD107a distribution. See Figure S3. **(B)** Histograms represent the mean and standard deviation of the percentage of asymmetric cells in 3 independent experiments. No statistical difference was revealed by paired t-test. **(C, D)** Statistical analysis of cells in telophase and in G1. (**C)** Cells in telophase are plotted against their CD107a FI. The different curves represent cells having undergone 0, 1 or 2 mitoses. Each dot indicates one cell undergoing asymmetric CD107a repartition as compared to its CD107a FI. The χ^2^ statistical test showed that cells undergoing uneven repartition of lytic machinery in telophase were randomly distributed all over the CD107a expression curves (See Materials and Methods section). **(D)** Plots show cells in G1 from three different experiments. Curves represent the distribution of CD107a florescence intensity for all cells in G1. Individual plots, marked with different colors, show cells in G1 at different rounds of division. The Kolmogorov-Smirnov goodness of fit test rejected the hypothesis that the CD107a expression curves follow the same distribution at the different division round (See Supplementary Results). The χ^2^ test showed that variability was distributed all over the curves. See Figure S3.

We next asked whether this process might create a drift in lytic compartment content in daughter cells leading to the emergence of cellular subsets expressing higher or lower levels of CD107a. To address this question, we analyzed the total CD107a-FI in all G1 cells (either undivided or following each division round). As shown in **Figure 5D**, the total CD107a-FI appeared to be broadly similar in the different rounds of division in the whole populations, suggesting that uneven repartition of CD107a, in a relatively constant fraction of cells at each division round, does not lead to the emergence of well-defined cellular subsets expressing higher or lower levels of CD107a. We employed the Kolmogorov-Smirnov goodness of fit test to determine whether the different curves followed the same distribution or not. The test strongly rejected the hypothesis that the CD107a expression curves follow the same distribution during the first two division rounds (see Materials and Methods section), indicating that during these division events randomly heterogeneous populations were generated. Nevertheless, our test also showed that the Kolmogorov distance decreased when the number of divisions increased, indicating that CD107a-FI density distribution seems to be convergent with a higher number of divisions. To define where variability was located in the curves, we employed the χ^2^ test. The test showed that variability was distributed all over the curves (i.e. for all the CD107a-FI). Together, Kolmogorov-Smirnov goodness of fit and χ^2^ tests revealed a non-stationary variability in the content of CD107a^+^ vesicles in CD8^+^ T cells during early division events. Taken together, the above results indicate that asymmetric distribution of CD107a^+^ vesicles in telophase is not limited to the first division, but it is rather a stochastic process, inherent to each division, that perpetuates variability in daughter cells.

### Lytic granules randomly distribute on the two sides of the cleavage furrow

To gain direct information about the possibility that lytic components might stochastically distribute in nascent daughter cells, we visualized lytic granule repartition during division in individual CTL transfected with mCherry-tagged GrzB mRNA, by live cell microscopy. mCherry-tagged GrzB showed no preferential localization within cell cytosol at the different phases of the division and appeared to randomly partition into the two nascent daughter cells. In some cases, nascent daughter cells exhibited approximately similar repartition of lytic granules (**Figure 6A, Video 2**), in some other cases lytic granule repartition appeared to be rather asymmetric (**Figure 6B, Video 3**). Furthermore, we investigated cell division in 4D (3D plus time). Sorted CD8^+^ T cells in G2/M phase were loaded with LysoTracker Red (LTR) to stain their late endosomal lysosomal vesicles (of which lytic granules are an important fraction (Faroudi et al., 2003)). Nascent daughter cells were imaged to monitor distribution of LTR^+^ vesicles and measure the integrated fluorescence intensity. An example of one CD8^+^ T cell distributing LTR^+^ vesicles in a symmetric fashion during division is shown in **Figure 6C** and **Video 4** (LTR distribution ranged within 40-60% at all time points measured). One CD8^+^ T cell that distributed in an asymmetry fashion LTR^+^ vesicles is shown in **Figure 6D** and **Video 5** (LTR distribution ranged above or below 40-60% at all time points measured). Additional examples of cells dividing in symmetric and asymmetric fashion are shown in **Figure 6-figure supplement 1** and **Video 6.**

**Figure 6:**
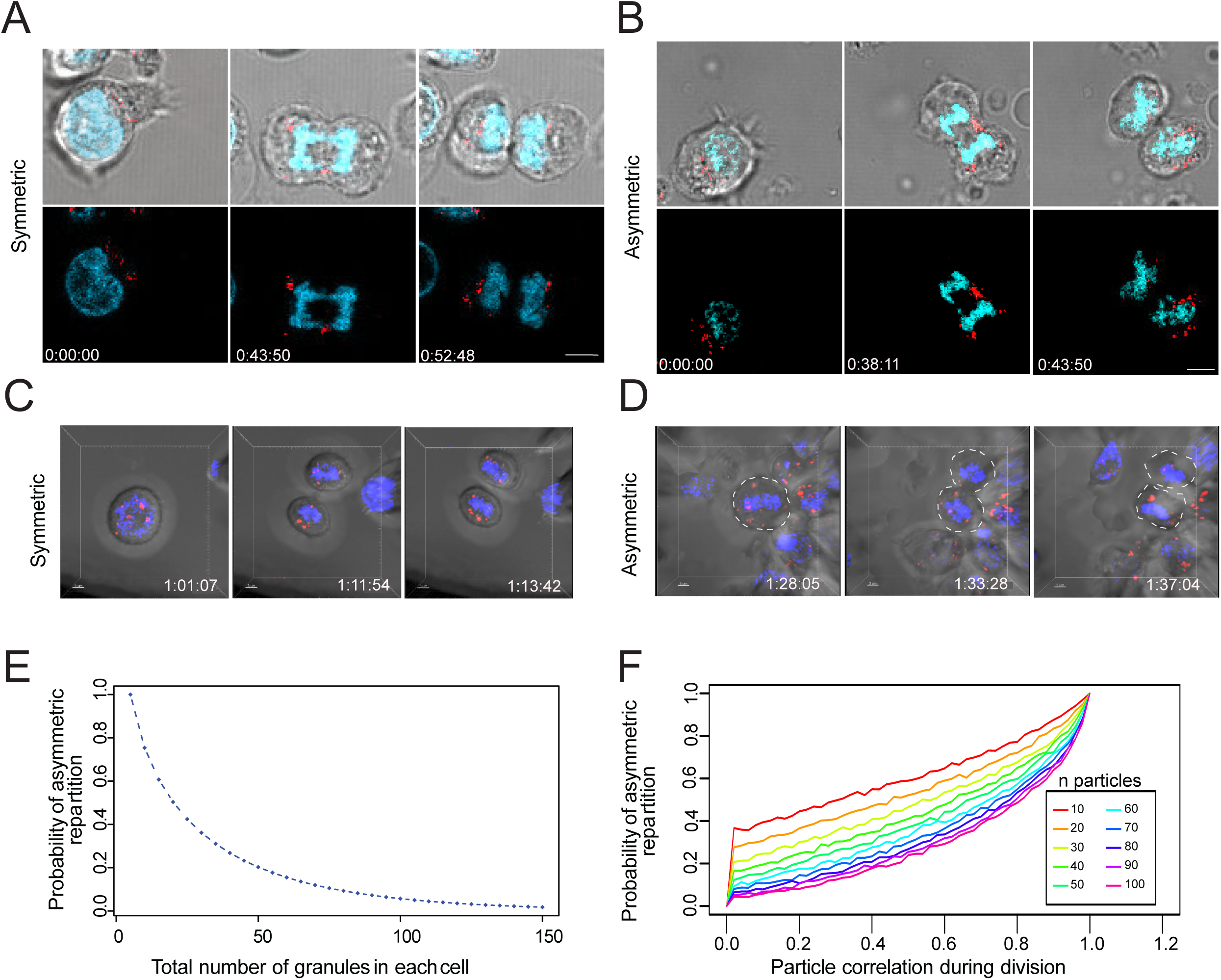
Lytic granules randomly distribute on the two sides of the cleavage furrow. **(A and B)** Snapshots depict typical cells in division undergoing even (**A**) or uneven (**B**) repartition of lytic granules (mCherry-tagged GrzB, red) in telophase as detected by live cell imaging. Images are from Video 2 and 3 respectively. Results are from 3 independent experiments. **(C and D)** Snapshots depict Imaris software reconstructions of typical cells undergoing even **(C)** or uneven **(D)** repartition of LTR^+^ (red) lytic granules in division as detected by 4D live cell imaging. Images are from Video 4 and 5 respectively. Results are from 4 independent experiments. See Videos 4-6. (**E**) Binomial modeling for the behavior of the population of n granules. The curve shows the probability of lytic granule asymmetric repartition in telophase as a function of lytic granule number. (**F**) Monte-Carlo simulation of particle correlation as a function of lytic granule number and probability of lytic granule asymmetric repartition.

While lytic granules seemed to be overall randomly distributed between nascent daughter cells, in some cases the videos showed that lytic granules did not behave completely independently from each other and exhibited some clustering. We therefore used a computational approach to establish whether the above-described process might be linked to a random repartition of lytic components into the two nascent daughter cells. We first calculated the probability to obtain an asymmetric distribution of lytic granules (*e.g.* a repartition of the granules into the two daughter cells out of the 40-60% range) related to the granule number per dividing cell. This computation is naturally handled with a binomial modeling for the behavior of the population of n granules (see Materials and Methods section). This analysis showed that for n <100 the probabilities that individual particles distribute asymmetrically on the two sides of the cleavage furrow are relatively high (**Figure 6E**). Using stimulated emission depletion (STED) on CTL stained for GrzB, we estimated that 14 to 65 (mean = 37) lytic granules are contained within individual CTL. Our estimation well-matched with numbers published in previous studies, ranging between 10 and 100 (Chiang et al., 2017; Clark et al., 2003; Kataoka et al., 1996; Peters et al., 1991). These values are compatible with a significant probability of stochastic uneven repartition of lytic granules in telophase, assuming that all lytic granules behave independently.

Since our videos indicate that lytic granules might form transitory aggregates within confined intracellular spaces, we upgraded our mathematical simulation of lytic granule repartition in telophase to include the possibility that lytic granules might not segregate completely independently. We simulated particle correlation during cell division for 10 to 100 particles. To evaluate the correlation level between individual particles during cell division (ranging from 0 = absence of correlation to 1 = 100% correlation) for a given probability of asymmetric repartition (outside the interval [40%-60%]), we used a Monte-Carlo approach (see Material and Methods section).

The analysis shows that for a probability of 20% asymmetric repartition of particles (corresponding to 20% uneven repartition of lytic granules during cell division experimentally measured by imaging flow cytometry and confocal imaging), particle correlation has a relatively low value (4% for 37 particles, 0.04; CI95%, 0.035-0.045), suggesting that lytic granules mainly segregate independently during cell division. Taken together, cell imaging and computational results strongly suggest that the observed stationary unequal distribution of lytic granules in telophase is the result of a stochastic repartition of particulate cytosolic structures on the two sides of the cleavage furrow in dividing cells.

### The level of lytic granule content in individual CTL dictates CTL killing capacity

To assess the consequences of an uneven distribution of lytic compartment on CTL-mediated cytotoxicity, we investigated cytotoxic efficacy in CTL expressing high and low lytic granule content. Clonal CTL were loaded with LysoTraker blue, and cells containing high (LysoTracker^High^) and low (LysoTracker^Low^) levels were FACS sorted. As shown in **Figure 7A**, sorted LysoTracker^High^ and LysoTracker^Low^ CTL populations maintained their difference in LysoTracker staining at least 24 hours after cell sorting. The cytotoxic efficacy of sorted CTL populations was compared at different effector:target (E:T) ratios by measuring the percentage of killed targets (7-AAD positive targets). For each ratio, LysoTracker^High^ CTL were more efficient than LysoTracker^Low^ CTL in exerting cytotoxicity (**Figure 7B-C**), although basal killing (in the absence of peptide stimulation) was comparable between LysoTracker^High^ and LysoTracker^Low^ CTL (**Figure 7C**). The above results indicated that lytic granule content is associated with killing efficacy. To strengthen these findings, we performed additional experiments on sorted CTL for high and low LysoTracker staining and measured CD107a surface exposure and CD8 internalization following 4 hour conjugation with target cells. Results show that LysoTracker^high^ CTL exhibited a higher lytic granule secretion as detected by CD107a exposure when compared to LysoTracker^low^ CTL (**Figure 7D**). However, productive TCR engagement was comparable in both populations as detected by similar levels of CD8 internalization (Huang et al., 2019; Xiao et al., 2007) (**Figure 7E**).

**Figure 7:**
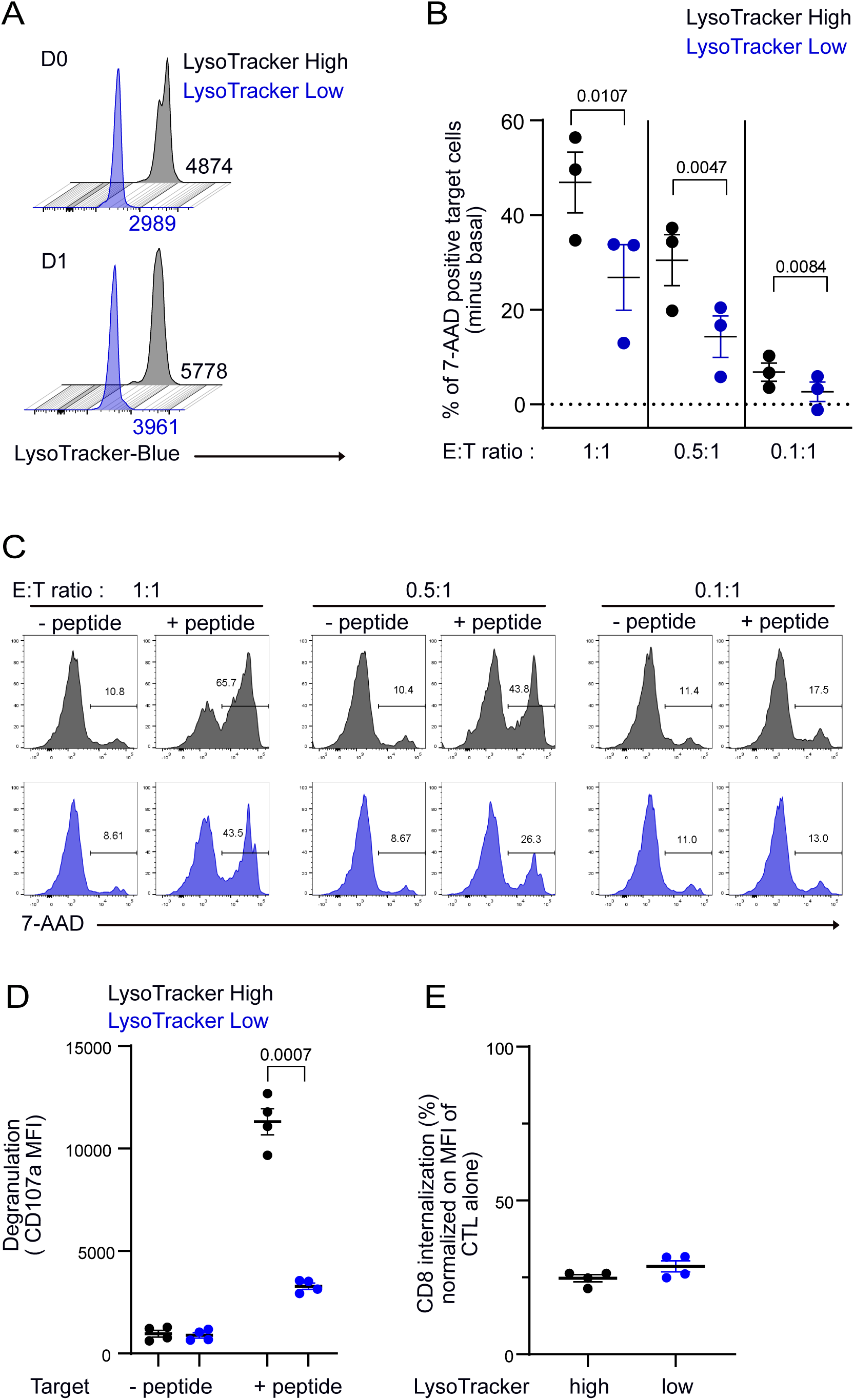
CTL expressing high level of lytic granules have better killing capability. Clonal CTL were FACS-sorted on the basis of their LysoTracker Blue staining. (**A**) Representative FACS histograms showing LysoTracker Blue staining levels on LysoTracker^high^ and LysoTracker^low^ sorted-CTL at the indicated day (D) after cell sorting. Numbers indicate mean fluorescence intensity. Results are representative of 3 independent experiments **(B-C)** LysoTracker^High^ and LysoTracker^Low^ CTL-mediated cytotoxicity was evaluated by FACS analysis by measuring 7-AAD uptake in target cells either pulsed or not with antigenic peptide following overnight incubation with CTL at the indicated E/T ratio. (**B**) Cytotoxicity is expressed as the % of 7-AAD^+^-pulsed target cells minus % of 7-AAD^+^-unpulsed target cells (basal). Results are from 3 independent experiments. Each dot represents results from one experiment performed in triplicate. Means +/- SEM are shown. Paired t-tests were performed and P-values are indicated. (**C**) Histograms shown are from one representative experiment. Numbers indicate the percentage of 7-AAD positive target cells. (**D**) LysoTracker^High^ and LysoTracker^Low^ CTL CD107a exposure after a 4 hours incubation with target cells pulsed or not with antigenic peptide (E/T ratio 0.5:1) was evaluated by FACS analysis. Each dot represents results from 4 independent experiments performed either in duplicate or triplicate. Means +/- SEM are shown. Paired t-tests were performed and P-values are indicated. (**E**) CD8 expression in LysoTracker^High^ and LysoTracker^Low^ CTL after a 4 hours incubation with target cells pulsed with antigenic peptide (E/T ratio : 0.5:1) was evaluated by FACS analysis. Results are normalized on CD8 MFI level of LysoTracker^High^ and LysoTracker^Low^ CTL cultured in the absence of target cells. Each dot represents results from 4 independent experiments performed either in duplicate or triplicate. Means +/- SEM are shown.

Together, these results indicate that the lytic granule cargo of individual CTL and not their activation properties directly impact killing behavior. They imply that stochastic uneven distribution of lytic granules in dividing CTL determine heterogeneous killing behavior at the single cell level.

## Discussion

In the present study we found that, in both freshly isolated peripheral blood CD8^+^ T cells and clonal CTL, ∼ 20 percent of telophasic cells undergoes asymmetric distribution of the lytic compartment into the two daughter cells. Our results establish that CD8^+^ killing capacity is associated to lytic compartment level and strongly suggest that uneven lytic machinery repartition produces CD8^+^ T cell populations with heterogeneous killing capacities.

We used imaging flow cytometry, a technique that combines the advantages of flow cytometry and microscopy and allows the detection and analysis of rare cells within whole cell populations on the basis of their morphological and staining characteristics (Basiji and O’Gorman, 2015; Doan et al., 2018; Hritzo et al., 2018). We thus acquired and analyzed a significant number of relatively rare events of T cell divisions by precisely identifying cells in telophase. The use of CTV distribution as a parameter of global protein repartition in telophase together with the acquisition of an important number of cell divisions strengthens our analysis. In addition, we investigated lytic granule repartition in dividing CD8^+^ T cells by 3D confocal laser scanning microscopy and 4D live cell imaging. These techniques allowed visualization of lytic granule repartition in telophase with a high time/space resolution and strengthened imaging flow cytometry data by providing unambiguous visualization of lytic granule partitioning.

Our results demonstrate that the uneven lytic machinery distribution is not related to ACD. In mouse T lymphocytes, ACD has been reported as a mechanism contributing to the generation of effector/memory daughter cells following division of an individual naive T cell in response to polarizing cues (Arsenio et al., 2015; Chang et al., 2007). Establishment of asymmetry has been associated to the uneven inheritance by daughter cells of transcription factors such as c-Myc and T-bet known for their role in the induction of metabolic reprogramming and in the acquisition of T cell effector function respectively (Chang et al., 2011; Verbist et al., 2016). Following the original observation of uneven repartition of proteasomes in dividing mouse CD4^+^ T cells leading to asymmetric degradation of T-bet in daughter cells (Chang et al., 2011), additional cellular effectors including metabolic and signaling pathways have been found to be implicated in fate determining ACD in mouse naive T lymphocytes (Kaminski et al., 2016; Pollizzi et al., 2016; Verbist et al., 2016). Our results, by showing that lytic granule repartition is not accompanied by a detectable asymmetric segregation of T-bet and c-Myc and does not require a polarity cue, suggest that the lytic machinery uneven distribution observed in human CD8^+^ T cells is not related to previously described ACD. Although we could not detect an asymmetric repartition of classical lineage-determining transcription factor, in our models, this observation does not exclude the possibility that ACD might play a role in the differentiation of human naive T cells into effector and memory subsets during initial antigen specific immune responses. It is therefore possible that the discrepancy between our results and previous studies on ACD in mouse T lymphocytes arises from the different nature of the cells involved in the study. It should also be noted that, beside ACD, other mechanisms can contribute to the generation of different T lymphocyte populations from naive lymphocytes and, more in general, can play a role in T lymphocyte heterogeneity. Alternative models postulate that lymphocyte differentiation might be achieved via the accumulation of progressive differences among daughter cells due to variation in the quantity of the inherited proteins (Buchholz et al., 2016; Cobbold et al., 2018; Gerlach et al., 2013; Girel and Crauste, 2019; Pham et al., 2014; Rohr et al., 2014; Schumacher et al., 2010).

A puzzling question is how asymmetric distribution of lytic components in telophase is generated. Our results provide a stepping-stone to answer this question. First, mathematical analysis of our imaging flow cytometry data provides an interpretation of our results that is compatible with a stochastic distribution of lytic components during cell division. On one hand, mathematical analysis shows that the process of asymmetric distribution is stationary in terms of the fraction of involved cells: e.g. occurs always on a similar percentage of cells, at each division round, in different experiments and following different stimuli. On the other hand, the heterogeneity process, although stationary is not hereditary: e.g. a daughter cell originating from a heterogeneous division has a constant stationary probability to produce a new uneven division. Second, live-cell imaging shows lytic granule distribution during mitosis. We did not observe any specific pattern of lytic granule repartition (polarization at the membrane or close to the cleavage furrow) before or during cell division. Instead, lytic compartments appeared randomly distributed in cell cytosol. Our observations are consistent with the mathematical modeling of intracellular vesicle distribution showing the high probability of an uneven distribution of a relatively small quantity of granules. In other words, pre-packaged molecular components within a few relatively big vesicles might have higher probability to be asymmetrically partitioned in telophase than molecular components dispersed throughout the cytosol.

Moreover, it should be noted, that our videos and results in Figure 6 suggest that, in a limited number of division events, granules might not segregate completely independently from each other, implying that some active mechanism of granule segregation might contribute, to a minor extent, to lytic granule repartition in telophase.

Together, our results point out a mechanism of heterogeneity generation that is for the most part stochastic and might be a general mechanism for generating heterogeneity in dividing cells. The possibility that particulate material is unevenly distributed in telophase into two nascent daughter cells has been proposed for other organelles and in other cellular systems (Bergeland et al., 2001; Carlton et al., 2020; Sanghavi et al., 2018). Indeed, in MDCK cells, microscopy analysis and mathematical modelling based on the laws of probability suggested that endosomes/lysosomes partitioning between daughter cells is stochastic (Bergeland et al, 2001). Others show that in telophasic cells, endosomal compartments are clustered at the cleavage furrow, suggesting that microtubules are involved in this process. However, no mechanism ensuring endosomal compartment anchorage to either spindle has been revealed, suggesting that this repartition is stochastic. Similarly, in Dictyostelium cells, it has been demonstrated that dynein and kinesin motors drive phagosomes segregation independently of each other and stochastically (Shanghavi et al 2018). To our knowledge, our present study is the first to relate a mechanism of a random segregation of organelles to functional heterogeneity of immune cells.

What could be the functional role of asymmetric molecular segregation during mitosis in human CD8^+^ T cells? We propose that a mechanism of asymmetric distribution in telophase (that is stationary at each division, but not inherited by daughter cells) can be instrumental to randomly generate short-lived CTL cohorts harboring functional heterogeneity while ensuring globally reproducible antigen specific CD8^+^ T cell responses.

This process might confer robustness to CTL responses through population averaging (Buchholz et al., 2016; Hodgkin et al., 2014).

It is interesting to note that our results present analogies with previously published data in which asymmetric segregation of internalized exogenous antigen was found to occur during B cell division (Thaunat et al., 2012). Together with this previous study, our results reveal an intriguing capacity of both T and B cells to stochastically distribute in telophase their acidic compartments: MHC Class II compartments for B cells and lytic granules for CD8^+^ T cells. Thus, stochastic distribution in telophase appears to be a major mechanism ensuring a high variability of both humoral and cellular adaptive immune responses during lymphocyte clonal expansion.

## Material and Methods

### Key resources table

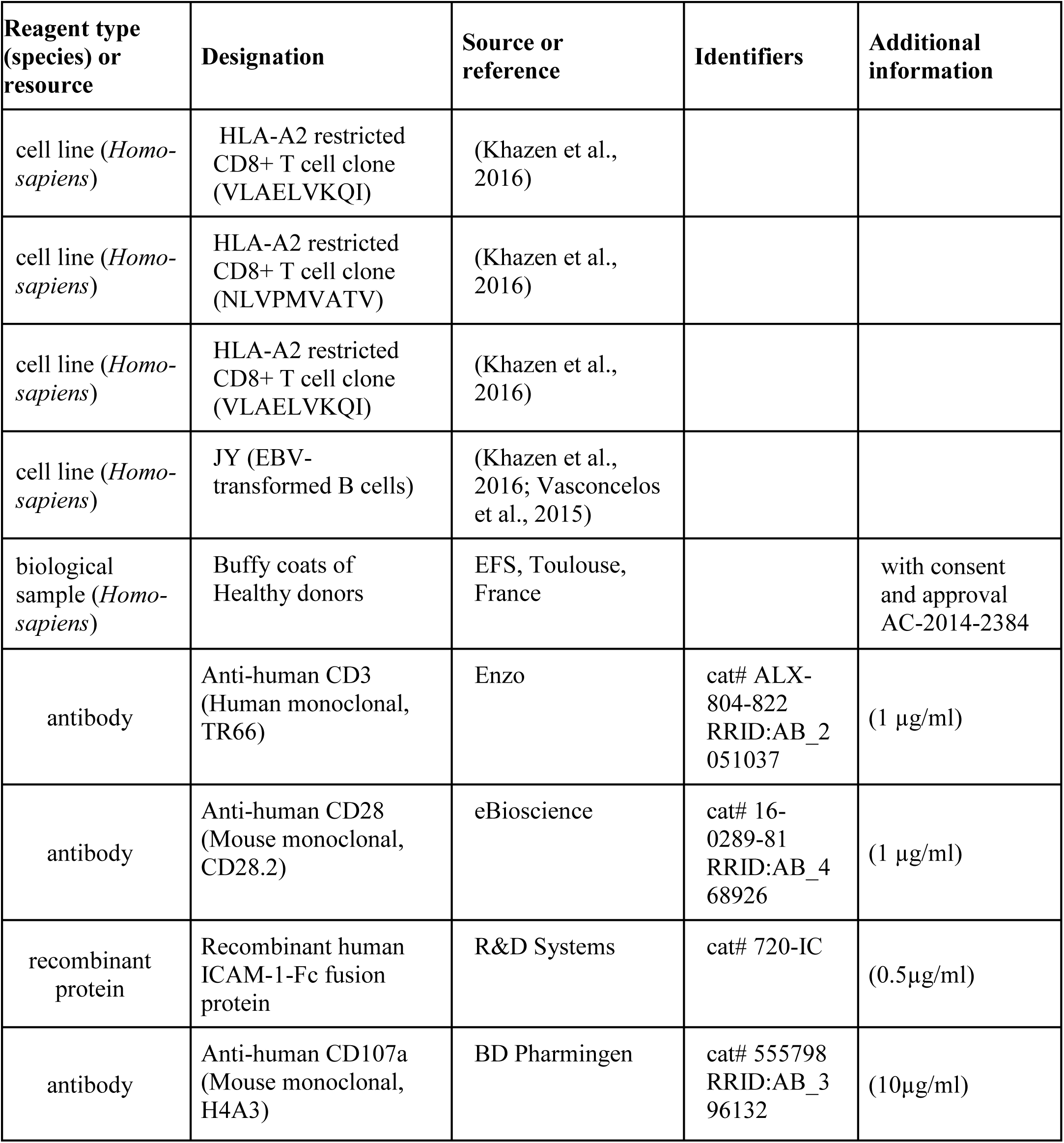
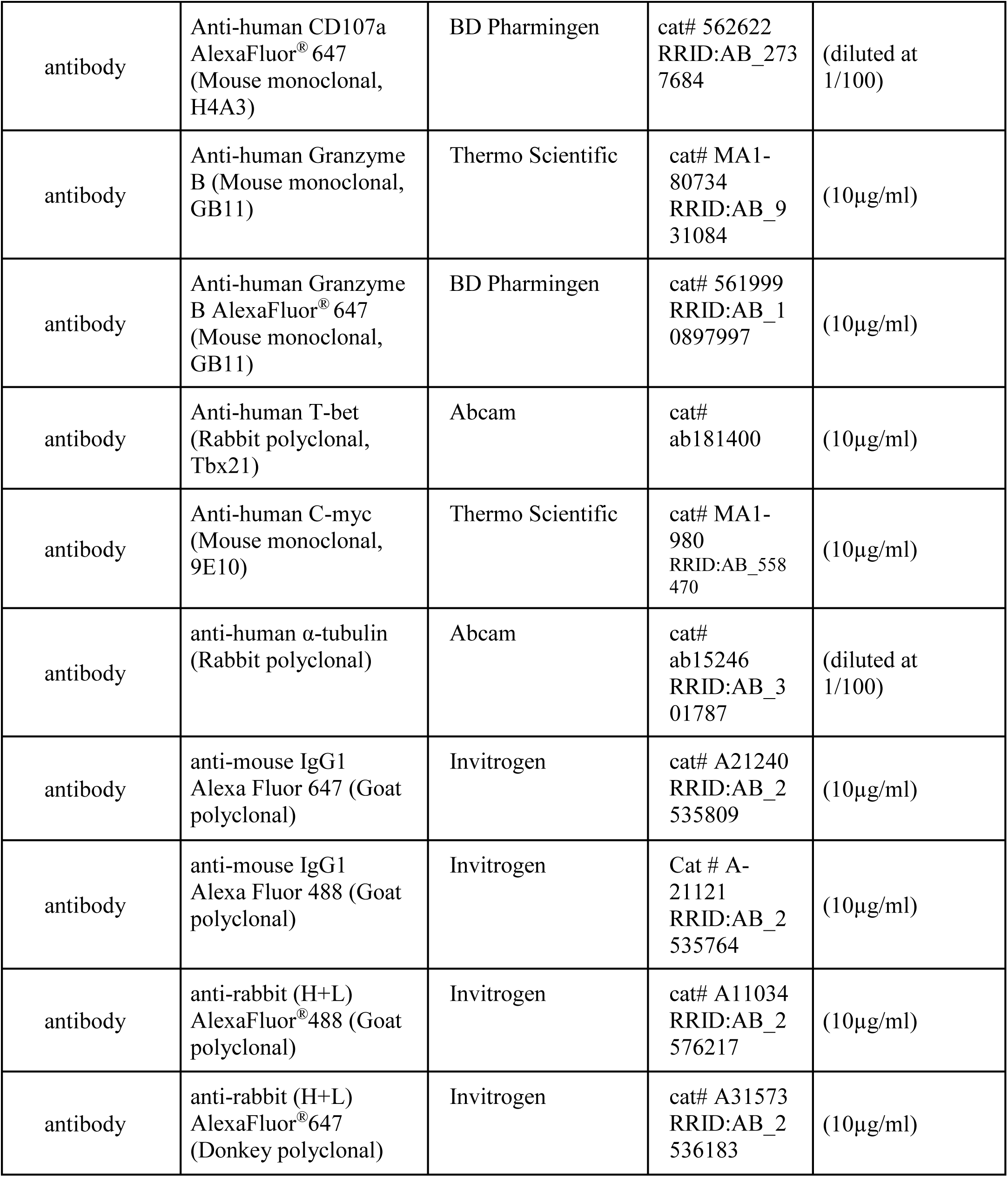
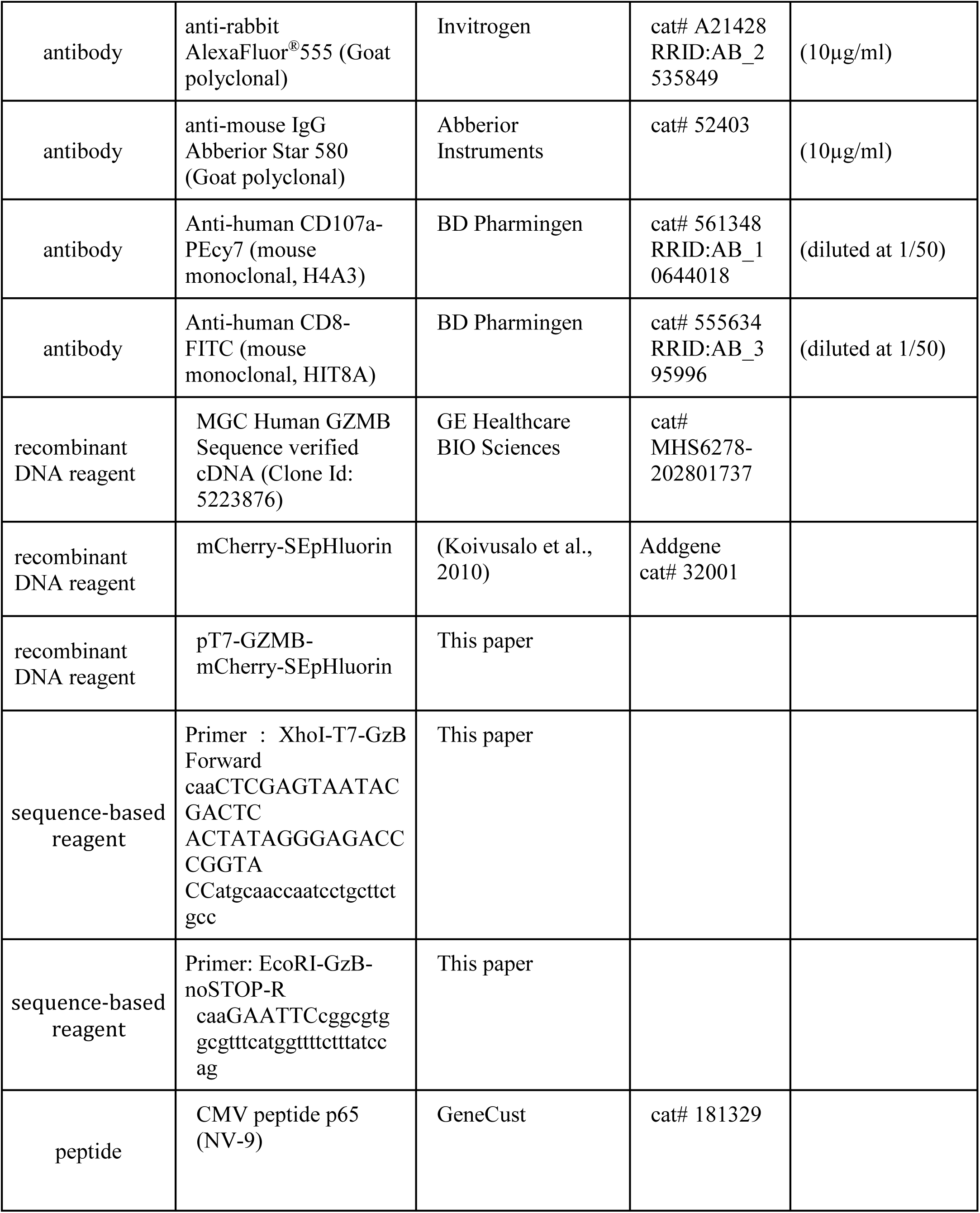
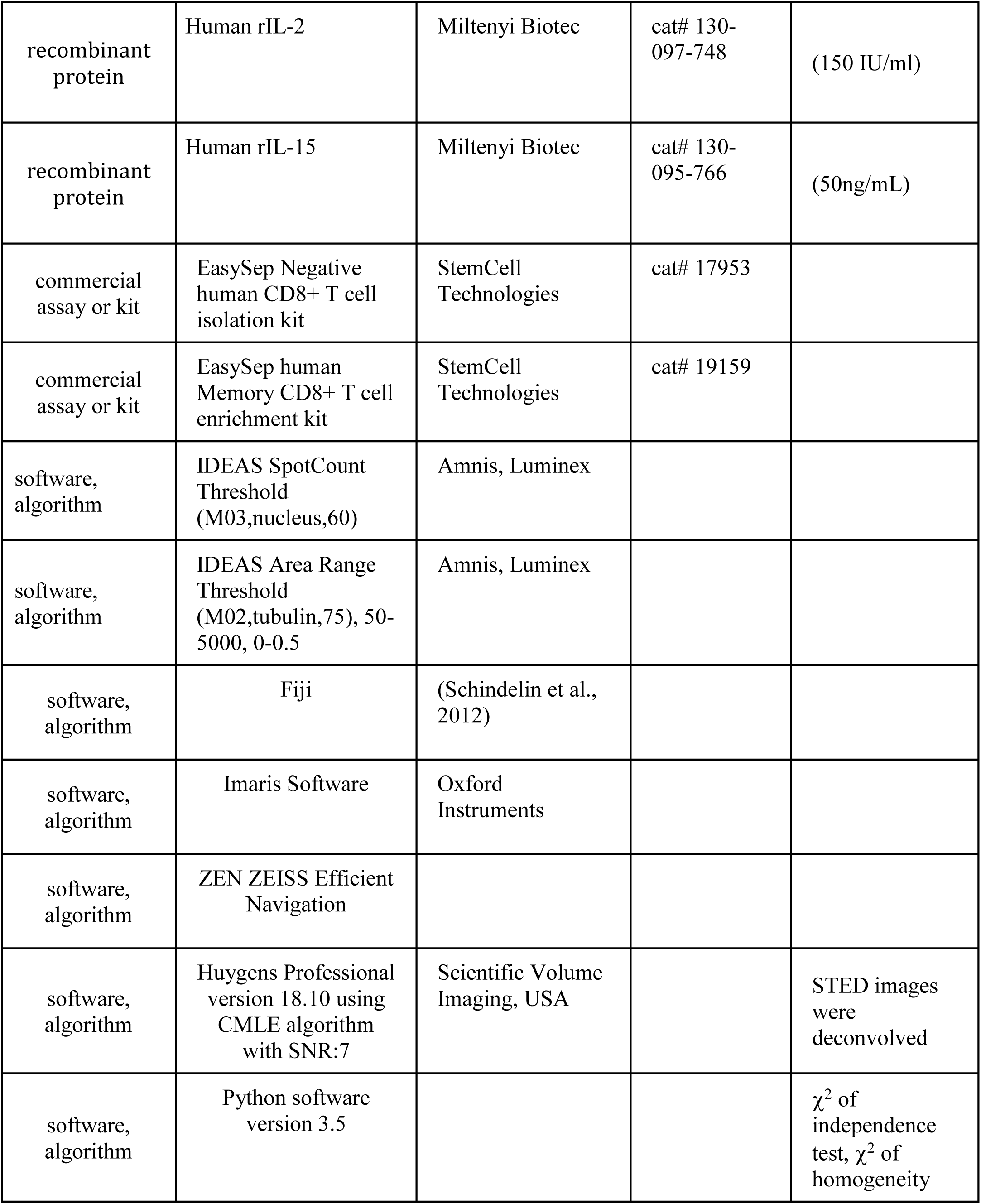
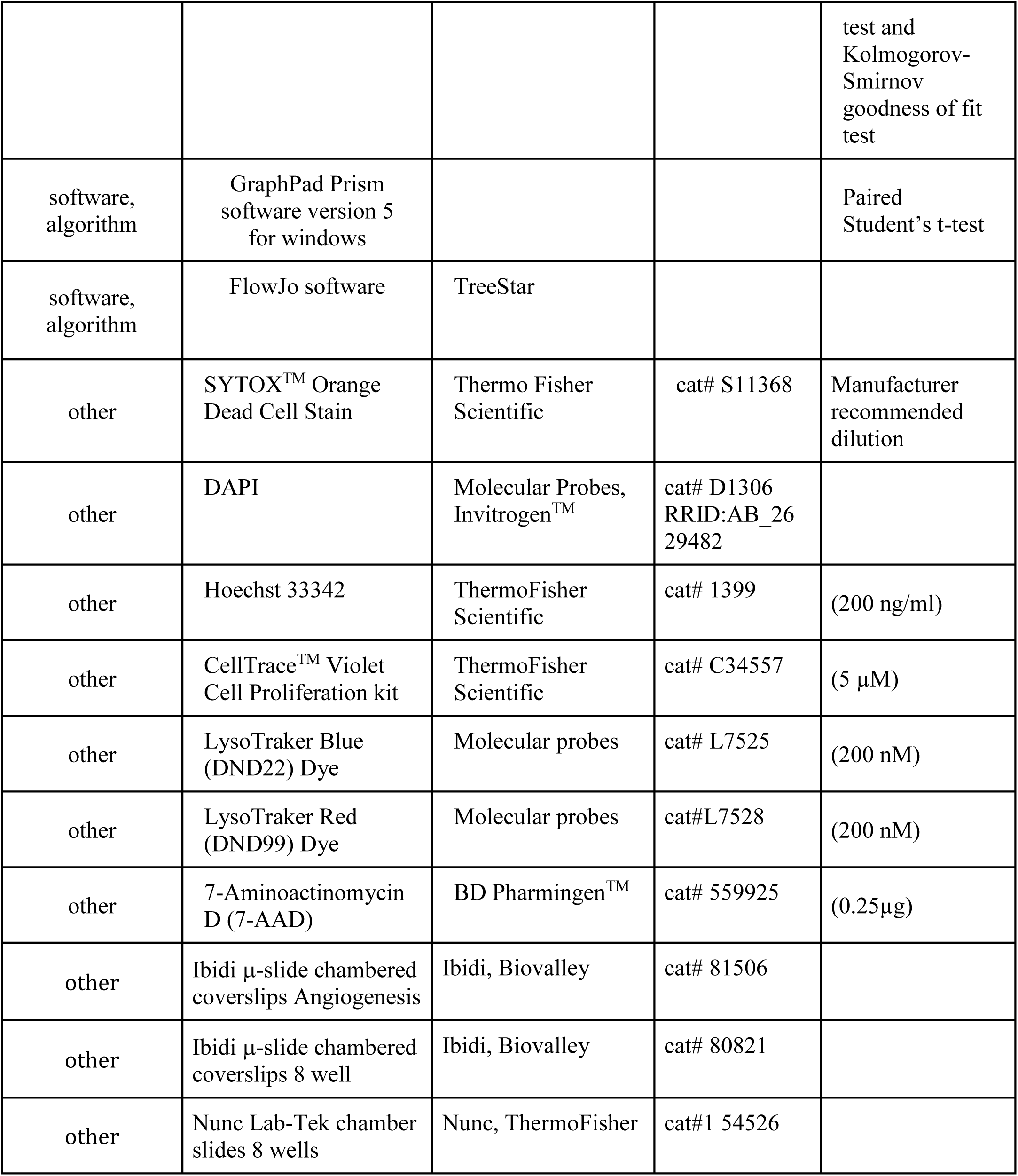
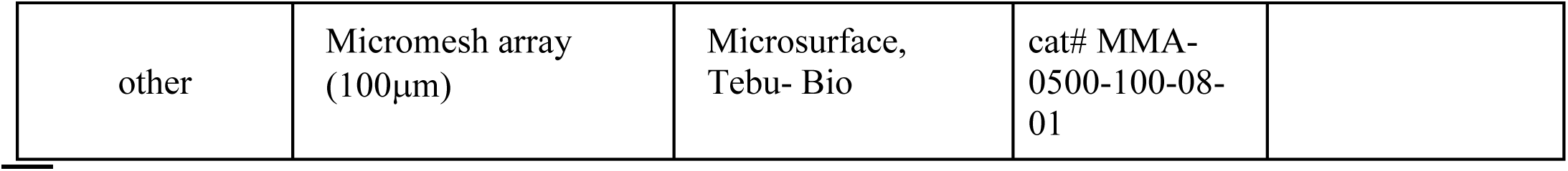

### Experimental model and subject details

Peripheral blood mononuclear cells were isolated from buffy coats of healthy donors obtained through the Etablissement Français du Sang (EFS, Toulouse, France). Blood samples were collected and processed following standard ethical procedures (Helsinki protocol), after obtaining written informed consent from each donor and approval by the French Ministry of the Research (transfer agreement AC-2014-2384). Approbation by the ethical department of the French Ministry of the Research for the preparation and conservation of cell lines and clones starting from healthy donor human blood samples has been obtained (authorization No DC-2018-3223).

### Cell culture and stimulating conditions

Total human CD8^+^ T cells were purified from healthy donor blood samples using the EasySep Negative human CD8^+^ T cell isolation kit (StemCell Technologies). CD8^+^ T cells were routinely ∼90% pure. Memory human CD8^+^ T cells were purified from healthy donor blood samples using the EasySep Human Memory CD8^+^ T cell enrichment kit (StemCell Technologies), cells were routinely ∼90% CD8^+^RO^+^RA^-^.

HLA-A2 restricted CD8+ T cell clones, specific for the NLVPMVATV peptide or the VLAELVKQI peptide of the CMV protein pp65 were cultured in complete RPMI/HS medium (RPMI 1640 medium supplemented with 5% human AB serum; Inst. Biotechnologies J. Boy, Reims), minimum essential amino acids, HEPES, sodium pyruvate (Invitrogen), 2-mercaptoethanol (5 µM, Gibco) and 150 IU/ml human rIL-2 and 50ng/ml rIL-15). Clones were re-stimulated every 2-3 weeks in 24-wells plate with 1×10^6^ irradiated (35 Gy) allogeneic peripheral blood mononuclear cells (isolated on Ficoll Paque Gradient from fresh heparinized blood samples of healthy donors, obtained from EFS) and 1×10^5^ irradiated EBV-transformed B cells. Complete RPMI/HS-Medium was supplemented with 1 µg/ml PHA.

EBV-transformed B cells (JY) HLA-A2+ were used as target cells and cultured in RPMI 1640 GlutaMAX supplemented with 10% FCS and 50 μM 2-mercaptoethanol, 10 mM HEPES, 1X MEM non-essential amino acids, 1X sodium pyruvate, 10 μg/mL ciprofloxacine. Profiling of JY cells has been done using STR.

All cell lines are routinely screened for mycoplasma contamination using MycoAlert mycoplasma detection kit (Lonza, Basel, SW).

For imaging flow cytometry (ImageStream®X, Merk) and confocal laser-scanning microscopy human CD8^+^ T cells or CD8^+^ T cell clones were stimulated for 72h with immobilized anti-CD3 (1 µg/ml, TR66, (Valitutti, 1995)), anti-CD28 (1 µg/ml, clone CD28.2, eBioscience) and immobilized recombinant ICAM1-Fc fusion protein (0.5µg/ml, R&D Systems) in complete RPMI/HS medium. For confocal laser-scanning, cells were plated on anti-CD3/CD28/ICAM1 coated Nunc Lab-Tek Chamber Slide™ system 8 wells at 500 000 cells / well. For image stream analysis, cells were plated on anti-CD3/CD28/ICAM1 coated 24 well plates at 1.5×10^6^ cells / well.

### Image Stream analysis

#### Staining and acquisition strateg

Cells were first stained with CellTrace ^TM^ Violet Cell Proliferation Kit (CTV) in PBS (5 µM, 20 min, 37°C). After 72 hours of stimulation (cf: Cell culture and stimulating condition), cells were fixed in 1% PFA (10 min, 37°C) and permeabilized in permeabilization buffer (PBS 3% BSA, 0.1% saponin, Sigma) for 5 min. Cells were incubated for 45 min with the indicated antibodies: AlexaFluor^®^ 647 anti-human CD107a antibody (diluted at 1/100, clone H4A3; BD Pharmingen ^TM^), anti-human Perforin (10µg/ml, clone δG9; BD Pharmingen ^TM^), AlexaFluor^®^ 647 anti-human Granzyme B antibody (10µg/ml, clone GB11, BD Pharmingen ^TM^), anti-human T-bet (Tbx21) (10µg/ml, clone 4B10; Abcam), anti-human C-myc (10µg/ml, clone 9E10; Thermo scientific), anti-human α-tubulin (diluted at 1/100, rabbit polyclonal; Abcam). The following secondary antibodies were used: AlexaFluor^®^488 or 647 goat anti-mouse IgG1 (10µg/ml; Invitrogen), AlexaFluor^®^488 or 647 anti-rabbit (H+L) (10µg/ml; Invitrogen). For image acquisition, cells were adjusted to 10.10^6^ -20.10^6^ /mL in FACS buffer (PBS, 1% FCS, 5% Hepes, 0.1% Azide) containing SYTOX^TM^ Orange Dead Cell Stain (recommended dilution, Thermo Fisher Scientific) for nuclear staining. Cells were acquired using ImageStream®X (IsX; Amnis, Luminex) technology.

#### Telophase discrimination strategy

Amnis IDEAS software was used to analyze IsX data and identify cells in telophase. As in classical cytometry data analysis, cells in G2/M phase were first selected according to their DNA content (fluorescence of SYTOX orange). A mask based on nuclear staining was employed (SpotCount Threshold (M03, nucleus, 60)) to visualize the nuclei of cells/events in the G2/M fraction at the single cell level. A second mask (Area Range (Threshold (M02, tubulin, 75), 50-5000, 0-0.5)) based on the α-tubulin staining (to clearly identify the narrow intracellular bridge of highly condensed α-tubulin that participates to midbody formation) was employed to distinguish telophases from anaphases or cell-doublets. Finally, the results from both masks were used to manually verify that selected cells were cells unambiguously in telophase.

#### Analysis of cell protein distribution during telophase using Fiji

Staining intensities of α-tubulin, CTV and of the different markers of interest were analyzed on Fiji to determine the percentage of proteins inherited by the two nascent daughter cells in telophase. Watershed function of Fiji software was used on the α-tubulin staining intensity to determine the specific areas corresponding to the two nascent daughter cells in telophase. The obtained areas were converted to masks that were next applied to measure CTV and the fluorescence of the different markers of interest. This procedure allowed us to determine the intensity of fluorescence in the two nascent daughter cells in telophase respectively. The percentage of staining in each nascent daughter cell was determined as: Fluorescence Intensity of daughter cell 1 / (Fluorescence Intensity of daughter cell 1 + Fluorescence Intensity of daughter cell 2) x 100. To test the specificity of the staining with the different antibodies used to study molecular repartition in telophase, we measured the fluorescent intensity of secondary antibodies or isotype controls as compared to specific antibodies. This analysis gave the following values of MFI: CD107a 70.527 isotype control 13.621; perforin 716.312, secondary mouse antibody 56.383; GrzB 677.445 isotype control 13.621; T-Bet 356.228 secondary mouse antibody 56.383; c-Myc 1.434.537 secondary rabbit antibody 14.231.

### 3D laser scanning microscopy on fixed cells

After 72 hours of stimulation, cells were fixed in 1% PFA (10 min, 37°C). Permeabilization and staining with antibodies were performed in PBS 3% BSA, 0.1% saponin (Sigma) for 5 min and 45 min respectively. The following antibodies were used: anti-human CD107a (10µg/ml, cloneH4A3, BD Pharmingen ^TM^) followed by AlexaFluor^®^488 goat anti-mouse IgG1 (10µg/ml; Invitrogen), anti-human α-tubulin (diluted at 1/100, rabbit polyclonal; Abcam) followed by AlexaFluor^®^555 goat anti-rabbit (10µg/ml; Invitrogen). Nuclei were labeled with DAPI (1µg/ml, 5 min). The samples were mounted in 90% glycerol-PBS containing 2.5% DABCO (Sigma) and examined using a LSM710 (Zeiss) confocal microscope with a ×63 plan-Apochromat objective (1.4 oil) with an electronic zoom of 4. Cells in telophase were identified on the basis of nuclear and tubulin marker staining. 3D images (using the z-stack function) were acquired for every cell identified as being in telophase. CD107a fluorescence intensity in the two nascent daughter cells was measured on 2-D image projections obtained applying the Sum function of Fiji Software to z-stack series. Since the background noise made the watershed function unsuitable to use, a region of interest (ROI) corresponding to the nascent daughter cell was manually drawn on the basis of brightfield and tubulin staining. We determined the percentage of CD107a staining in each nascent daughter cell as: CD107a intensity of daughter cell 1 / (CD107a intensity of daughter cell 1 + CD107a intensity of daughter cell 1) × 100.

### Stimulated Emission Depletion Microscopy

CTL were seeded on poly-L-lysin coated high performance coverslips and fixed in 3% PFA (10 min, 37°C). Permeabilization and staining were performed in PBS 3% BSA, 0.1% saponin (Sigma) for 5 min and 60 min respectively. Cells were stained with an anti-human Granzyme B antibody (10µg/ml, clone GB11, Thermo Scientific) followed by a goat anti-mouse IgG Abberior Star 580 (Abberior Instruments). Coverslips (high performance D=0.17mm +/-0.005, ZEISS, Germany) were mounted on microscopy slides using Mowiol-DABCO.

STED images were acquired with a Leica SP8 STED 3X microscope (Leica Microsystems, Germany) using a HC PL APO CS2 100X/1.4 oil immersion objective. To optimize resolution without bleaching in 3-D, the 775 nm STED lasers line was applied at the lowest power that can provide sufficient improvement in resolution compared to confocal. Z-stack series were acquired sequentially with the pulsed 532 nm laser. For image acquisition, we used the following parameters: 3 time average/line, 400 Hz scan speed. STED images were subsequently deconvoluted with Huygens Professional (SVI, USA) using the CMLE algorithm, with a signal to noise ratio (SNR) of 7. 3-D image visualization was performed using the Fiji software.

### Live cell imaging

For 3D live cell imaging, the T7 GZMB sequence was obtained by PCR amplification as a XhoI-EcoRI fragment from pCMV-SPORT6-GZMB by using XhoI-T7-GZB forward primer and EcoRI-GRZB noSTOP reverse primer (Employed primers: Name: XhoI-T7-GzB F caaCTCGAGTAATACGACTCACTATAGGGAGACCCGGTACCatgcaaccaatcctgcttctgcc Name: EcoRI-GzB-noSTOP-R caaGAATTCcggcgtggcgtttcatggttttctttatccag).

XhoI-EcoRI fragment was cloned as a mCherry-SEpHlurin fusion construct in the pmCherry-SEpHlurin vector to produce the vector pGZMB-mCherry-SEpHluorin available to *in vitro T7 transcription. The plasmid* pCMV-SPORT6-GZMB and pmCherry-SEpHlurin were purchased from Addgene.

For efficient transfection of human CTL with tagged molecules allowing to monitor lytic granule repartition during mitosis, we synthetized capped and tailed poly(A) mCherry-tagged Granzyme B mRNA by *in vitro* transcription from the plasmid pGZMB-mCherry-SEpHluorin. One microgramme of pGZMB-mCherry-SEpHluorin was first linearized by NotI digestion to be used as templates for *in vitro* transcription by the T7 RNA polymerase using mMESSAGE mMACHINE T7 Ultra kit as per manufacturer’s protocol.

Human CTL were transfected using a GenePulser Xcell electroporation system (BioRad). 1x10^6^ CTL (5days after restimulation therefore in expansion phase) were washed and resuspended in 100μl Opti-MEM medium (Gibco) at RT with 2μg mCherry-tagged Granzyme B mRNA (*square wave* electrical pulse at 300V, 2ms, 1 pulse). 18 hours after transfection the transfection efficacy was verified by FACS analysis (typically 50-80%). Transfected CTL were seeded into poly-D-lysine-coated eight-well chambered slides (Ibidi, Munich, Germany) before imaging. Chambered slides were mounted on a heated stage within a temperature-controlled chamber maintained at 37°C and constant CO_2_ concentrations (5%) and inspected by time-lapse laser scanning confocal microscopy (LSM880, Zeiss, Germany with 1 image /30 seconds) for additional 5-6 hours using a Tile Scan mode to enlarge the acquisition fields and capture the rare cells undergoing spontaneous division during the time of acquisition.

For 4D live cell imaging, 72 hours after stimulation, CD8^+^ T cells were stained with Hoechst (200 ng/ml, ThermoFisher Scientific) to sort cells in G2/M phase by flow-cytometry (BD FACSAria SORP, BD Biosciences). Sorted cells were stained with LysoTracker Red (200 nM ThermoFisher) for 30 min at 37°C and washed. 20 000 cells in 5% HS/IL2/IL15 complete RPMI medium supplemented with 10 mM HEPES were seeded into poly-D-lysine-coated eight-well chambered slides (Ibidi, Munich, Germany) pre-coated with PDMS micromesh arrays (Microsurfaces, Melburn, Australia) containing 100-μm-diameter wells. Cells were 4D imaged (time and z-stack) on a heated stage within a temperature-controlled chamber maintained at 37°C and constant CO_2_ concentrations (5%) and inspected over night by time-lapse laser scanning confocal microscopy with a Plan-Apochromat 40x/1.3 Oil DIC M27 using an LSM780 or LSM880, Zeiss, Germany) or by spinning disk time-lapse microscopy using a spinning-disk microscope (Nikon) running on Metamorph software. A camera emCCD Evolve (Photometrics) was used for acquisitions. Image analysis was performed using Fiji software and 4-D videos and snapshots were generated with Imaris software.

### Cytotoxicity assay

CTL were incubated with 200nM LysoTracker Blue^®^ a probe staining the acidic lytic compartment of these cells (Faroudi et al., 2003) for 30 minutes at 37°C/5% CO_2_ in 5% FCS/RPMI/HEPES. After washing, cells expressing the highest and lowest 5-10 % LysoTracker Blue staining were sorted using a FACSARIA-SORP (BD Biosciences). CTL were used for standard over-night killing assays on the day of cell. Target cells were left unpulsed or pulsed with 10µM antigenic peptide during 2 hours at 37°C/5% CO2, washed three times and subsequently transferred to a 96 well U-bottom plate at 10×10^3^ cells/100μl RPMI, 5% FCS/HEPES. CTL were added to the target cells at the indicated effector (CTL): target (JY) ratio, in 100μl RPMI, 5% FCS/HEPES. Cells were pelleted for 1 min at 455 g and incubated at 37°C/5% CO2 overnight. Before FACS analysis, 0.25µg 7-Aminoactinomycin D (7-AAD; BD Biosciences) and FITC conjugated anti-CD8 antibody were added to each sample in order to measure the percentage of dead target cells. For the CD107a exposure and CD8 internalization assay, sorted CTL were incubated with target cells at 0.5:1 E/T ratio for 4 hours. Cells were stained with PE-cy7 conjugated anti-CD107a antibody and FITC conjugated anti-CD8 antibody for 30 min in FACS-buffer (1% human serum, 1% Foetal Calf Serum in PBS), washed, acquired on a Fortessa flow cytometer (BD Biosciences) and analysed by using FlowJo software (TreeStar).

### Statistical methods

*Paired Student’s t-test,* was performed to determine the statistical significance of differences between the groups (GraphPad Prism software version 5).

*Chi-square of independence test* was performed to determine the independence between the level of expression of a given marker and the capacity of a cell in telophase to asymmetrically distribute this marker (Python software version 3.5).

*Kolmogorov-Smirnov goodness of fit test* was performed to compare law between probability distribution of a marker of interest in cells in G1 (Python software version 3.5). *Chi-square of homogeneity test* was performed (in addition Kolmogorov-Smirnov goodness of fit test) to determine where the probability distribution of a marker of interest varies (Python software version 3.5).

### Statistical procedures

In the independence Chi-square test, we compare the theoretical effective (*e_i,j_*) to the observed effective (*n_i,j_*). The test statistic is defined by:

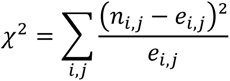

We compare it to 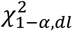, the quantile of the *χ*^2^ distribution associated to the 1 − α quantile. The quantile with 1 − α = 95% is the value such that 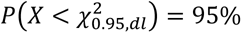 where P stands for the probability distribution of the Chi-square statistics with the associated degree of freedom dl.

We reject the hypothesis of independence between division of heterogeneous cells and division of all cells in one experiment when 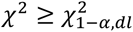 or when the p-value p satisfies *p* < *α* = 5%.

The red boxes represent the situations where we do not reject the hypothesis of independence of division between heterogeneous cells and all cells in one experiment. We shall observe that we never reject the hypothesis of independence.

The Kolmogorov-Smirnov test consists in analyzing if two independent samples follow the same law comparing their cumulative distribution function. We denote the two samples *X*_1_, *X*_2_, … *X_m_* and *Y*_1_, *Y*_2_, … *Y_m_* If we denote by *F_n_* and *F_m_* their cumulative distribution respectively, the test statistic is defined by:

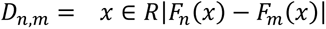

We compare it to *d_n,m,1_* _− *α*_, the quantile of the associated Kolmogorov-Smirnov distribution.

We then reject the hypothesis of adequation between cells of one division and cells of one other division in one experiment when *D_n,m_*≥ *d_n,m,1_* _− *α*_ or when the p-value p satisfies *p* < *α* = 5%.

The red boxes represent the situation where we do not reject the hypothesis of adequation between cells in one division and cells in another division. The white box represents the situation where we reject this hypothesis.

## Probability of an asymmetric repartition of lytic granules

To obtain a tractable formula for the computation of the probability of an asymmetric repartition of lytic granules, we use a binomial model that translates that each granule possesses a probability of 0.5 to attain each of the two daughter cells. The binomial model also assumes that all the granules behave independently of each other.

In that case, the probability of an asymmetric division for n granules is then equal to

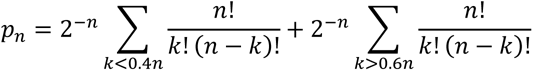

To evaluate the correlation level between particles (between 0 and 1) for a given probability of asymmetric repartition (outside the interval [40%-60%]), we use a Monte-Carlo approach where we sampled a sequence of correlated random variables distributed according to a Bernoulli distribution of parameter 0.5 since to the best of our knowledge there is no explicit formula to calculate a such probability of asymmetric repartition. Even with a Monte-Carlo approach, the simulation is a little bit involved: if r is the correlation level and if *X_i_* is the value of the random variable at step i, then *X_i_*_+1_ is obtained by :

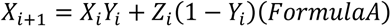

where *Z_i_* is a Bernoulli distribution of parameter 0.5 and *Y_i_* a Bernoulli distribution of parameter r. We shall verify that when *X*_1_, *X*_2_, … *X_n_* are sampled according to Formula A, they are Bernoulli distributed and pairwise correlated with a correlation r. Hence, we then mimic the correlated division with this model and then estimate the probability of asymmetric repartition with 5000 Monte-Carlo simulations for each value of r and a size of n=90 cells. We then evaluate the desired probability for r varying in a regularly spaced grid from 0 to 1 with a space equal to 0.02.

## Acknowledgements

We thank Dr. Stephane Manenti for discussion and critical reading of the manuscript ; Dr. Hellen Robey and Dr. Pauline Gonnord for discussion. We thank Dr. Liza Filali for advice in image analysis and Dr. Juliet Foote for critical reading of the manuscript. We thank the flow cytometry and imaging core facilities of the INSERM UMR 1043, CPTP and of the INSERM UMR 1037, CRCT, Toulouse, France.

This work was supported by grants from the Laboratoire d’Excellence Toulouse Cancer (TOUCAN) (contract ANR11-LABEX), Region Occitanie (contracts RCLE R14007BB, 671 34 No 12052802, and RBIO R15070BB, No 14054342), Fondation Toulouse Cancer Santé (contract 2014CS044) from the Ligue Nationale contre le Cancer (Equipe labellisée 2018) and from Bristol-Myers Squibb (No CA184-575). R.J. was supported by the Ligue Nationale contre le Cancer.

The funders had no role in study design, data collection and analysis, decision to publish, or preparation of the manuscript.

## Supplementary figure legends

**Figure 1-supplement 1.**
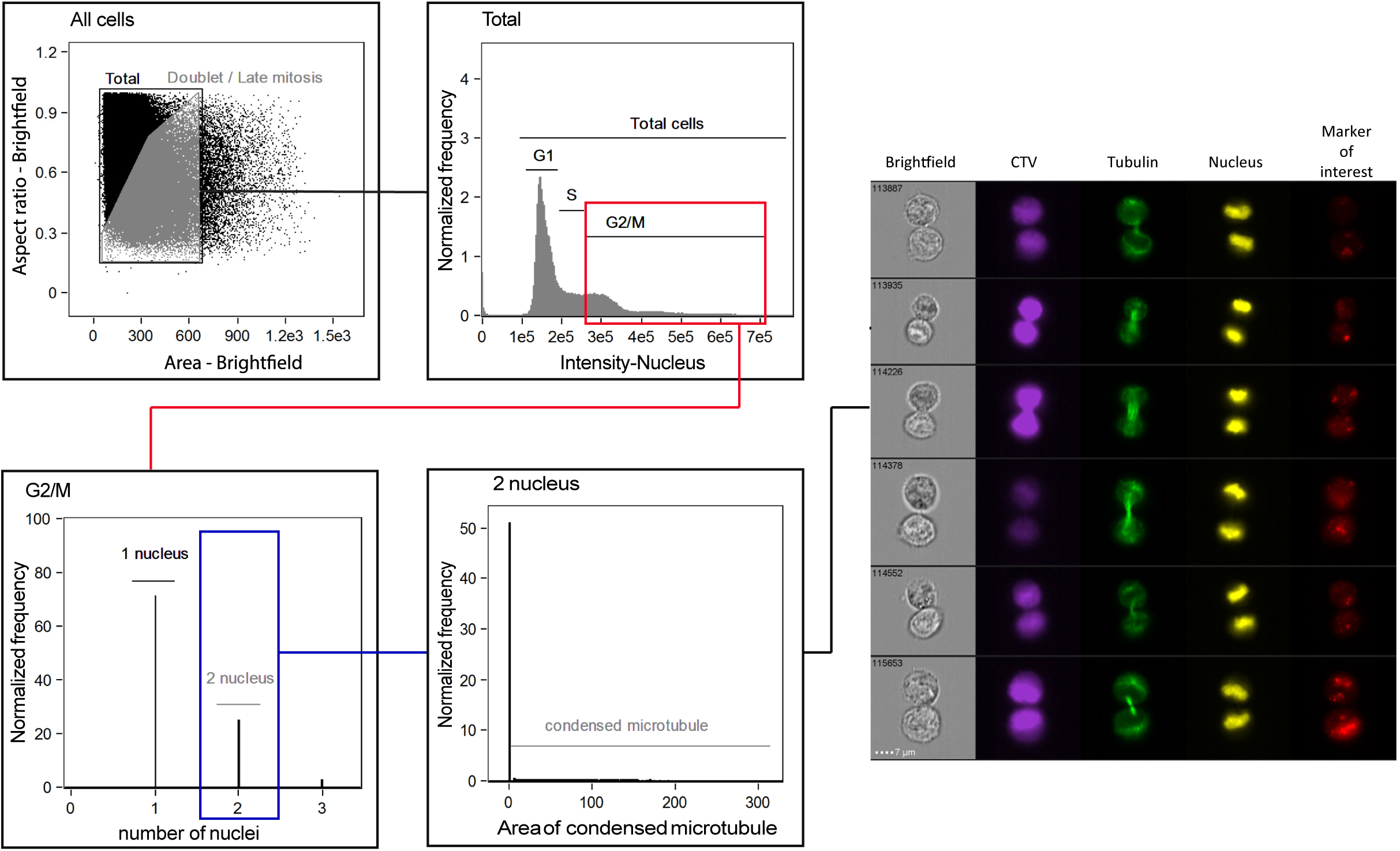
Gating strategy for Imaging Flow Cytometry (IsX) acquisition. Based on the brightfield illumination, all events were plotted for their aspect ratio (length/width, equal 1 for perfectly round cells) and their area. Cells in telophase were defined as those exhibiting a low aspect ratio and a big area. The region of interest (gray) included cell doublets and cells in anaphase and telophase. Based on the intensity of DNA staining (represented in linear axis) cells in G2/M were selected. We then applied a mask on the IsX image gallery (as described in material and methods section) to define the limits of the nuclei. This strategy was used to determine the number of nuclei present in each gated cell. To unambiguously identify cells in telophase we applied a mask on α-tubulin staining allowing to detect condensed microtubules in an elongated shape (as described in material and methods section). This procedure allowed us to detect the midbody (a structure characteristic of telophase formed by highly condensed α-tubulin that bridges the 2 nascent daughter cells). Cells included the described gates were finally visually inspected. All the cells recognized as in telophases on the basis of nuclear and tubulin staining were included in the analysis of the markers of interest.

**Figure 1-supplement 2.**
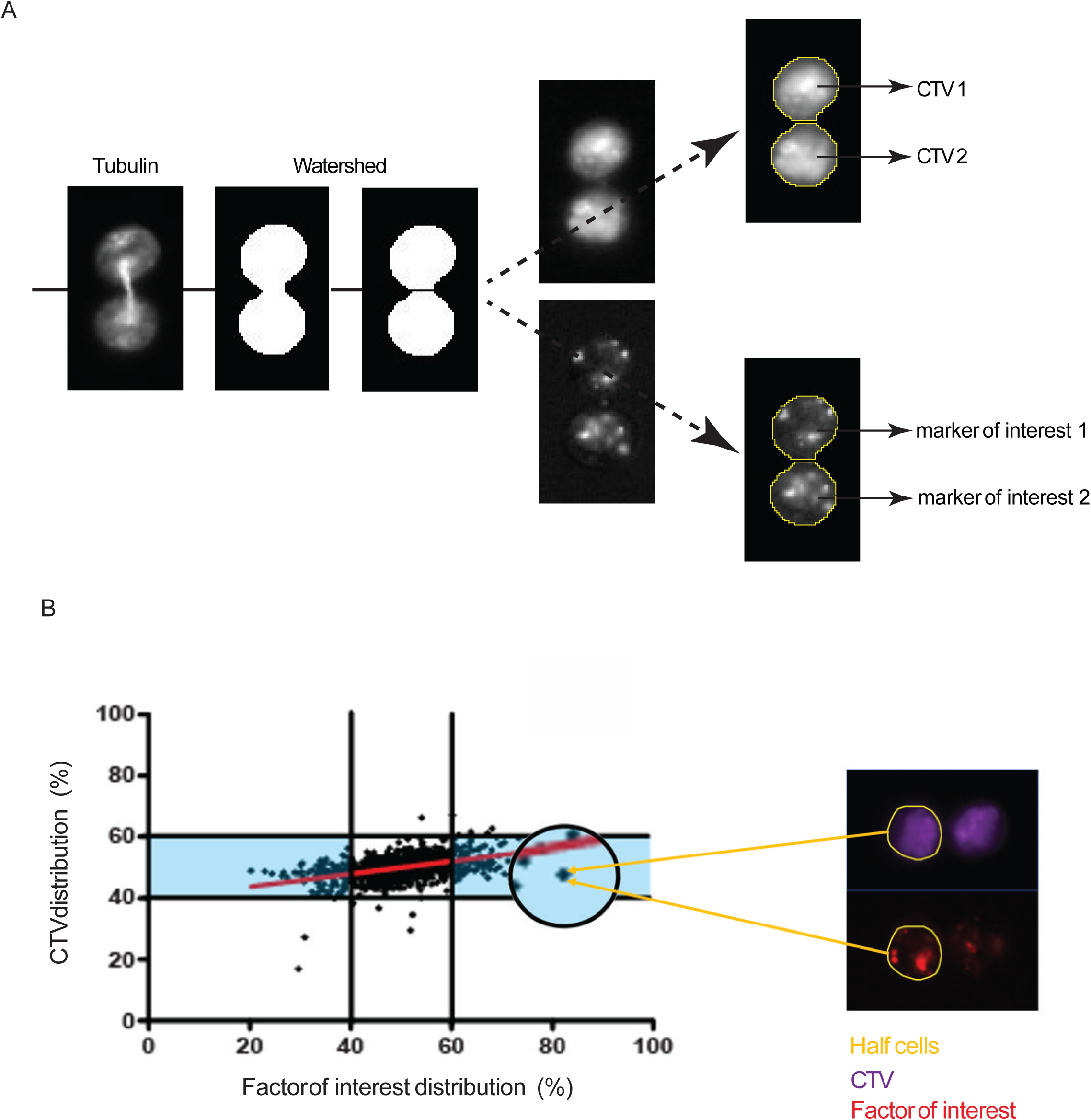
Analysis and representation of the repartition of markers of interest in dividing cells. **(A)** Analysis of individual cells in telophase. All IsX generated TIFF files were analyzed using the Fiji software. For each telophase cell we used 3 TIFF image corresponding to: i) CTV staining; ii) α-tubulin staining iii) and marker of interest. To standardize analysis, we used macro programming on Fiji (described in supplementary results section). To determine a rupture zone between the 2 nascent daughter cells we applied watershed function on tubulin mask. The watershed masks were used to determine the 2 nascent daughter cells in which the fluorescence intensities of CTV and of the markers of interest were measured (yellow lines). **(B)** Example of a cell exhibiting asymmetric distribution in telophase of a marker of interest. The yellow lines highlight the nascent daughter cell exhibiting a higher content of the marker of interest.

**Figure 1-supplement 3.**
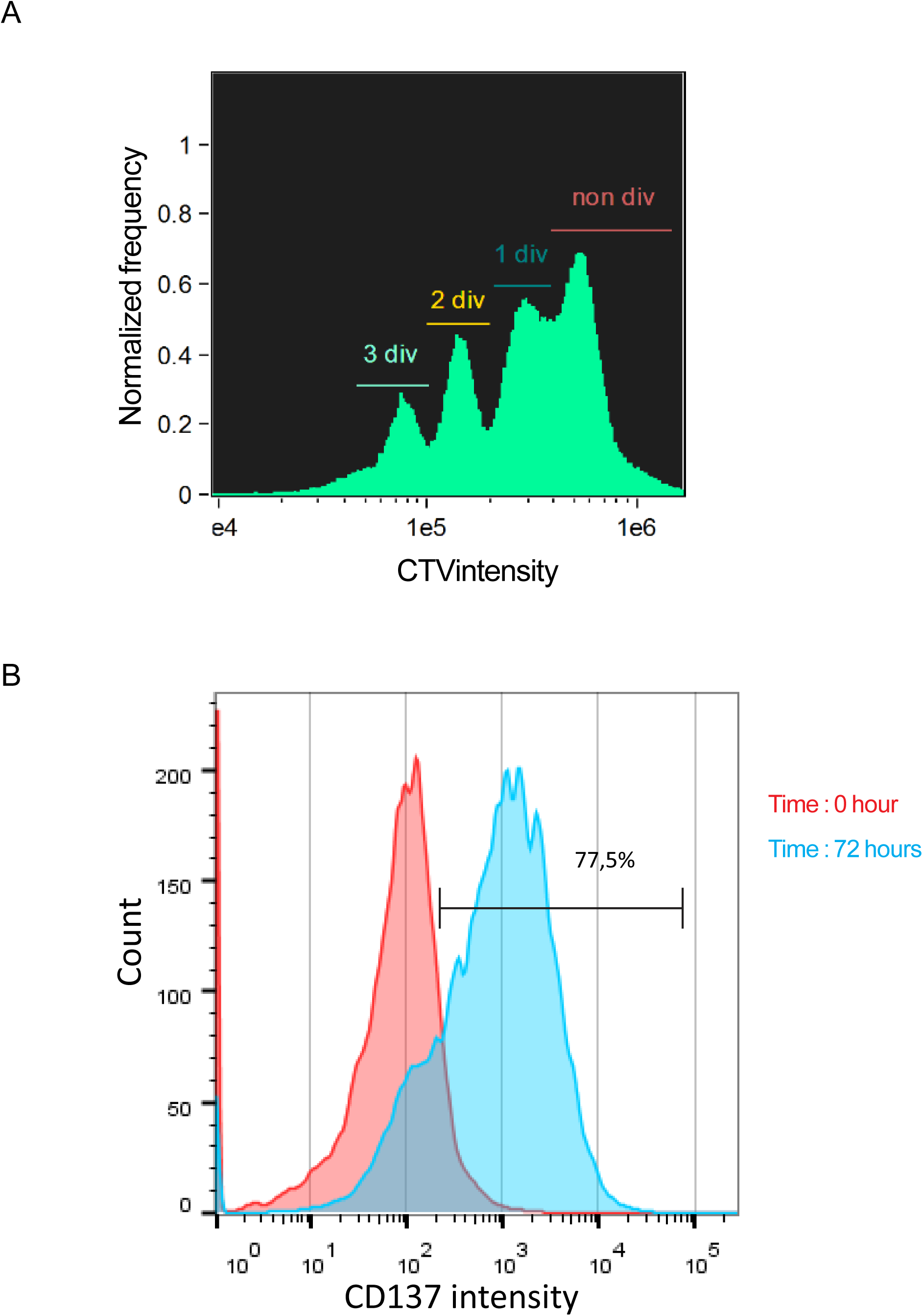
CD8^+^ T cells are efficiently stimulated on coated anti-CD3/anti-CD28/ICAM1. Freshly isolated polyclonal CD8^+^ T cells previously stained with CTV were stimulated 72 hours using immobilized anti-CD3/anti-CD28/ICAM1. (**A**) Imaging Flow Cytometry shows that stimulated cell undergo several rounds of division as shown by CTV staining dilution. (**B**) Flow cytometry shows upregulation of CD137 expression in stimulated cells.

**Figure 1-supplement 4.**
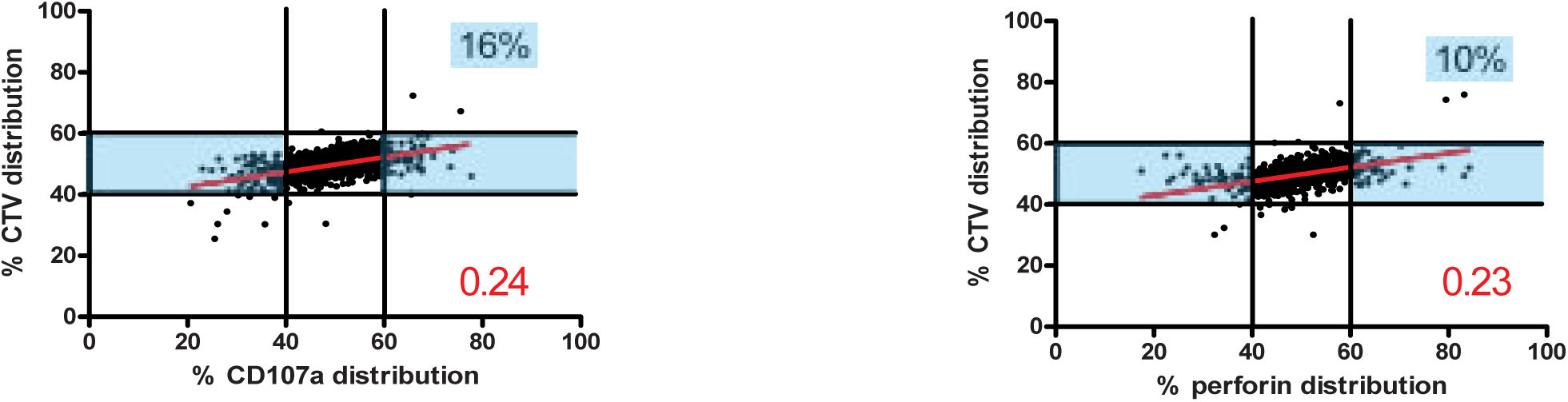
Uneven lytic granule segregation in telophase in CD8^+^ memory T cells. The panels show staining for CD107a and perforin in human CD8^+^ memory T cells stimulated and analyzed as **in Figure 1A-B**. CD107a n=978 from 3 independent experiments; perforin n=1127 from 3 independent experiments. Numbers highlighted in blue in the plots indicate the % of cells exhibiting asymmetric repartition of the marker of interest. Red lines indicate the global distribution of the data. Red numbers indicate the slope of the linear regression curve for marker distribution.

**Figure 6-supplement 1.**
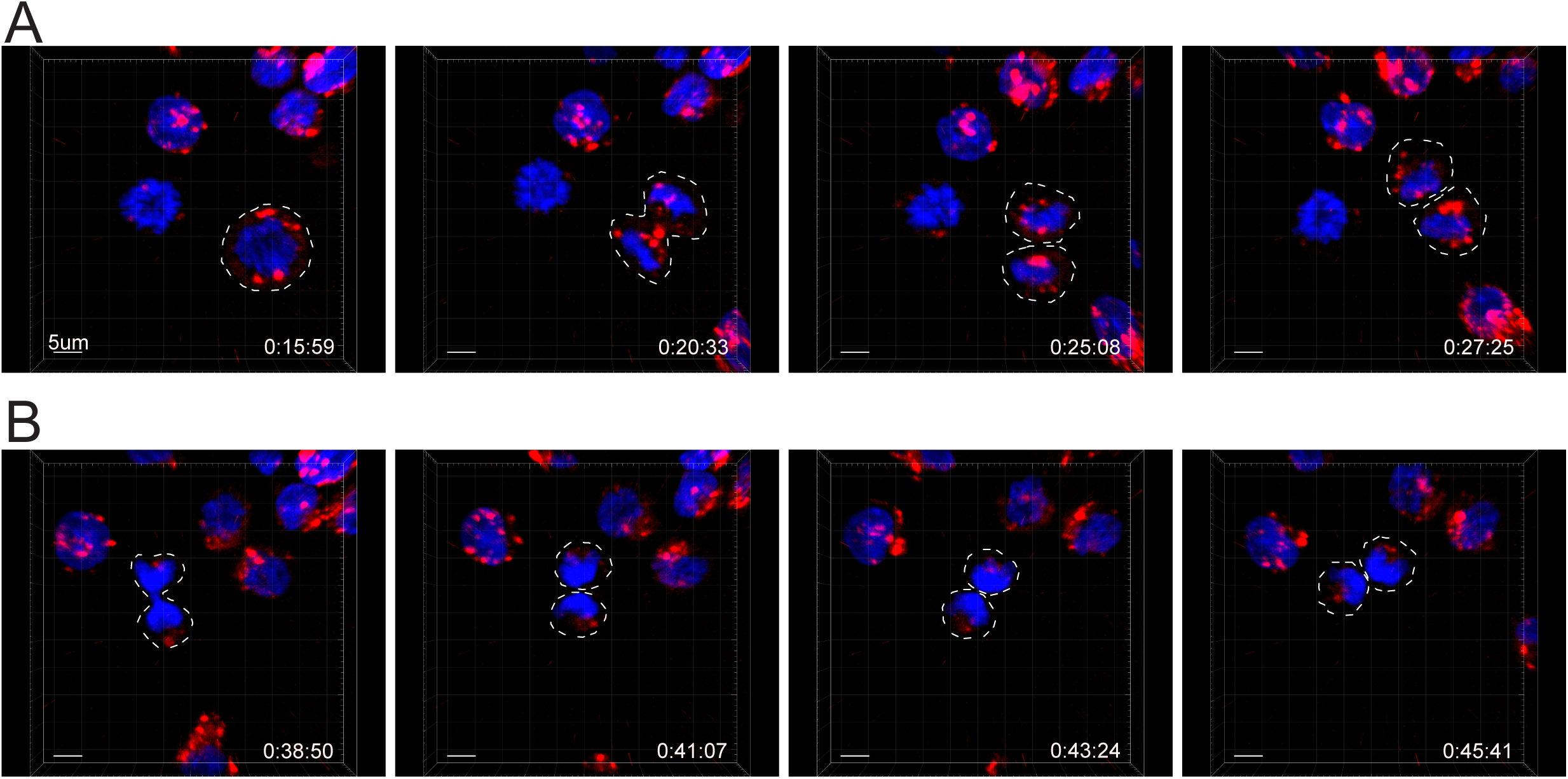
Lysotracker randomly distribute on the two sides of the cleavage furrow. **(A and B)** Snapshots depict Imaris software reconstructions of typical cells undergoing uneven **(A)** and even **(B)** repartition of LTR^+^ (red) lytic granules in division as detected by 4D live cell imaging. Images are from Video 6.

## Video legends

**Video 1: 3D visualization of CD107a repartition in a telaphasic CD8^+^ T cell** The video shows 3D reconstruction of a cell in telophase. CD107a (green), α-tubulin (red) and DAPI (cyan). The images presented in **Figure 2B** has been extracted from this video.

**Video 2 and 3: Visualization by time-lapse confocal laser scanning microscopy of cell division.** Human CTL were transfected during their expansion phase with mcherryGrzB. 18 hours after transfection, cells were inspected by time-lapse laser scanning confocal microscopy for additional 5-6 hours using a Tile Scan mode to enlarge the acquisition filed and to capture rare cells undergoing spontaneous division during the time of acquisition. Video 2 shows a typical cell undergoing even repartition of GrzB^+^ granules. Video 3 shows a typical cell undergoing uneven repartition of GrzB^+^ granules. Snapshots of Video 2 and 3 are shown in **Figure 6 A** and **B.**

**Video 4-6: Visualization by 4D time-lapse microscopy of cell division.** G2/M sorted CTL were loaded with Hoechst (blue) and LysoTracker Red (LTR, red) and inspected by time-lapse laser scanning confocal microscopy (video 4-5) or spinning-disk microscopy (video 6) for 12h-16h. **Video 4** shows 4D reconstruction (using Imaris software) of a typical cell undergoing even repartition of LTR^+^ granules. **Video 5** shows 4D reconstruction (using Imaris software) of a typical cell undergoing uneven repartition of LTR^+^ granules. Snapshots of Video 4 and 5 are shown in **Figure 6 C** and **D. Video 6** shows 4D reconstruction (using Imaris software) of one typical cell undergoing uneven repartition of LTR^+^ granules and one typical cell undergoing even repartition of LTR^+^ granules. Snapshots of Video 6 are shown in **Figure 6-figure supplement 1.** Results are from 4 independent experiments.

